# Tracking transcription-translation coupling in real-time

**DOI:** 10.1101/2023.12.07.570708

**Authors:** Nusrat Shahin Qureshi, Olivier Duss

## Abstract

A central question in biology is how macromolecular machines function cooperatively. In bacteria, transcription and translation occur in the same cellular compartment and can be physically and functionally coupled. While several recently published high-resolution structures of the ribosome-RNA polymerase (RNAP) complex provided first mechanistic insight into the coupling process, we do not know how these structural snapshots are placed along a dynamic reaction trajectory. Here, we reconstitute a complete active transcription-translation system and develop multi-color single-molecule fluorescence microscopy experiments to directly and simultaneously track transcription elongation, translation elongation and the physical and functional coupling between the ribosome and the RNAP in real-time. Our data show that the ribosome slows down while colliding with the RNAP and that coupling following a collision becomes less efficient. Unexpectedly, physical coupling can occur with hundreds of nucleotides of intervening mRNA between both machineries by mRNA looping, and increases in efficiency in presence of NusG. We detect active transcription elongation during mRNA looping and show that NusA-paused RNAPs can be activated by the ribosome by long-range physical coupling. We provide an alternative explanation on how the ribosome can rescue RNAP from frequent pausing without requiring collisions by a closely trailing ribosome. Our data mechanistically highlight an example of how macromolecular machines central to gene expression physically and functionally cooperate.

## Introduction

A central but understudied question in biology is how different macromolecular processes functionally cooperate. In bacteria, transcription performed by RNA polymerase (RNAP) and translation performed by the ribosome occur in the same cellular compartment, which allows functional coupling between both processes^1,2^. Such coupling provides opportunities for regulation that have been shown to make transcription more efficient in presence of active translation. For example, a ribosome closely trailing behind the RNAP inhibits Rho-dependent transcription termination by blocking the access to the Rho helicase^3^. In vitro, a closely trailing ribosome can also prevent RNAP pausing by suppressing hairpin-stabilized pause formation, or rescue paused RNAPs from backtracking by ribosome/RNAP collisions^4-6^.

Transcription and translation rates are also known to be correlated^6^, raising the question how elongation speeds in both processes are communicated between the two macromolecular machines. One possibility is that communication happens by the alarmone (p)ppGpp, which affects both transcription and translation elongation speeds^7^. Alternatively, direct physical interactions between the RNAP and a closely trailing ribosome could allow for coordinating elongation rates^4,5,8^. Indeed, several studies, including recent high-resolution structures, demonstrate a physical link that establishes coupling between both machines (termed expressome in this study) that is either mediated by direct interaction between the RNAP and the ribosome or by factor-mediated interaction mainly via the transcription factor NusG and/or NusA^5,9-13^. Those structures broadly fall into one of three categories: collided, coupled and uncoupled. The collided state, which contains the shortest possible intervening mRNA spacer between ribosome and RNAP, can neither bind NusA nor allow NusG-based bridging, potentially lacks the ω subunit of the RNAP and does not allow 30S head swiveling that is needed for ribosome translocation^10,11^. Further, it also could not be captured in vivo under normal growth conditions and could only be isolated when the cells were treated with the transcription inhibitor pseudouridimycin, artificially stalling the RNAP^12^. This raises the question whether ribosome/RNAP collisions, whereby the ribosome pushes the RNAP forward as recently suggested^4,5,13^, provide the prevalent mechanism for reactivating paused RNAPs in vivo. In contrast, the coupled state is compatible with known functional aspects of transcription and translation and NusA/NusG bridging, and therefore could provide a functional state during which communication between both machines could occur. However, the cryogenic electron microscopy (cryoEM) and cryogenic electron tomography (cryoET) structures only provide structural snapshots of a dynamic process and do not allow direct monitoring of activity. They also do not provide information on whether physical coupling can only occur when the two machines are close along the mRNA sequence or whether coupling could also occur via mRNA looping.

Here, we reconstitute the complete active transcription-translation system in vitro. We investigate the molecular mechanism of transcription-translation coupling by combining bulk biochemical experiments with multi-color single-molecule fluorescence assays to place various recently published structural snapshots along a reaction path. By simultaneously tracking transcription and translation elongation and the physical and functional coupling between both machineries, we find that the ribosome slows down before colliding with the RNAP, providing a biophysical explanation why the collided state is not found in vivo. We see that physical coupling can occur during active transcription elongation, by looping-out of the mRNA. This physical coupling allows the ribosome to activate a stalled RNAP by long-range communication, without the need for collisions, and provides an explanation how the central machineries in gene expression coordinate their activities.

### In vitro reconstitution of a fully active transcription-translation system

In order to directly visualize the process of transcription-translation coupling in real-time, we reconstituted the complete active transcription-translation system in vitro. We established single-molecule fluorescence microscopy assays that allow us to simultaneously track transcription and translation elongation occurring on the same nascent mRNA. To this end, we prepared a stalled transcription elongation complex (TEC) (Fig. 1a; see methods for details) consisting of a 3’-2xCy3.5 labeled DNA template, *E. Coli* RNAP and a nascently transcribed mRNA containing a ribosome binding site (RBS)^14^. We then loaded a Cy3-labeled 30S ribosomal subunit (labeled at an extension on h44 of the 16S rRNA) in presence of IF2a and fmet-tRNA^fmet^ on to the nascent mRNA^15,16^. The resulting translation pre-initiation complex (PIC) was immobilized via the 5’-end of the nascent mRNA to a PEG-biotin-functionalized glass surface for single-molecule multi-color imaging.

**Fig. 1:**
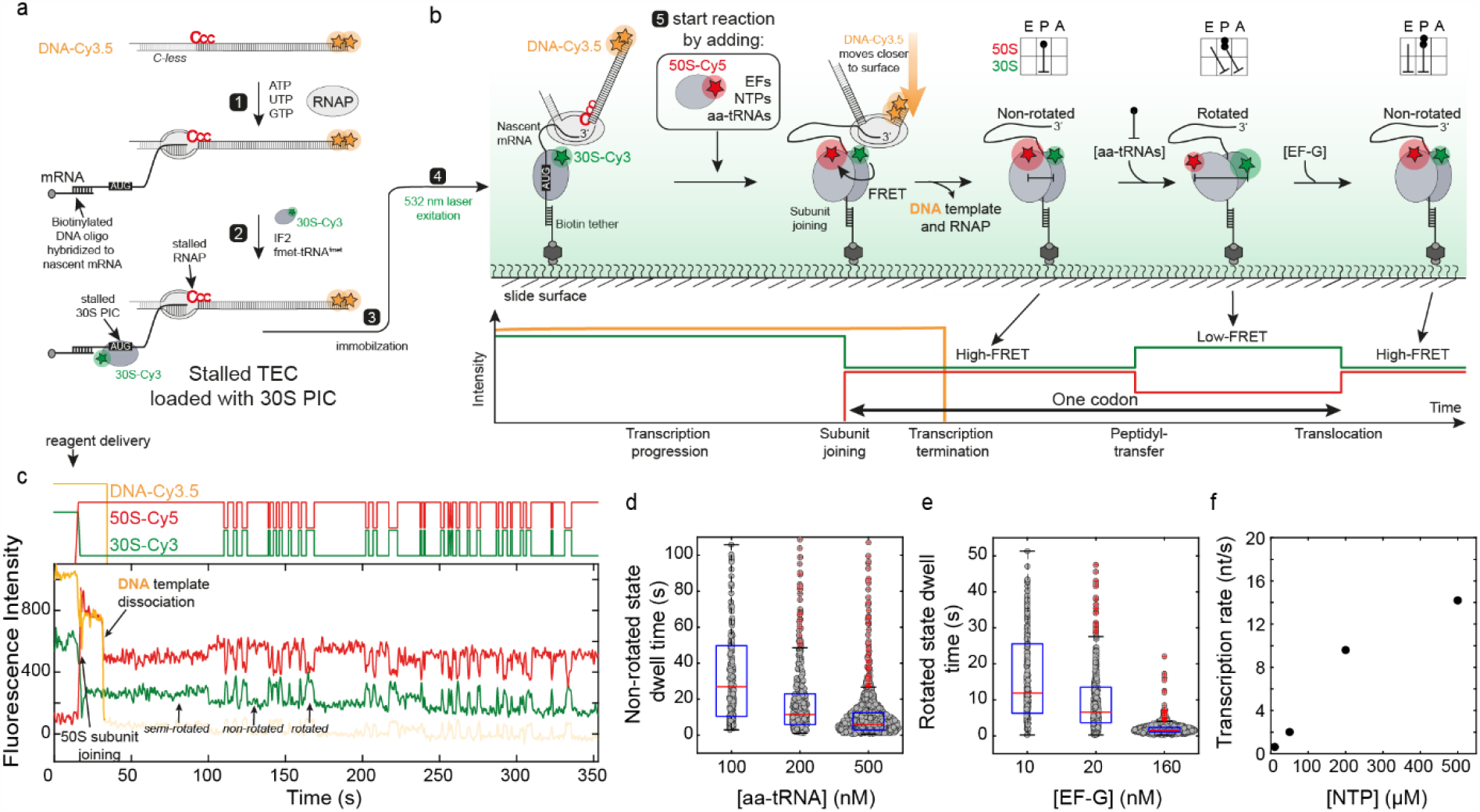
Single-molecule assay for real-time tracking of co-transcriptional translation elongation. **a**, Experimental design for preparation of transcription-translation complexes. **b**, Single-molecule assay to observe transcription and translation of the same mRNA molecules in real-time. **c**, Representative single-molecule trace showing fluorescence intensities of transcription (TC, yellow-orange) and translation (TL, red and green). Idealized trace is displayed on top. **d, e**, Beeswarm plot of dwell times for aa-tRNA-dependent non-rotated state (**d**) and EF-G-dependent rotated state (**e**). Median (red central mark), 25^th^ (bottom edge) and 75^th^ (top edge) percentile are indicated. Outliers are shown as red cross. Note that at the given y-axis range not all outliers are visible. Number of evaluated molecules: n=46 for 10 nM EF-G/100 nM aa-tRNA; n=53 for 20 nM EF-G/200 nM aa-tRNA; n=61 for 160 nM EF-G/500 nM aa-tRNA **f**, NTP-dependent average transcription rate (nt/s). Number of evaluated molecules: n=82 for 10 μM NTP; n=75 for 50 μM NTP; n=338 for 200 μM NTP; n=265 for 500 μM NTP.

Elongation of both machines was triggered at the beginning of imaging by delivering NTPs, Cy5-labeled 50S subunit (labeled at an extension on h101 of the 23S rRNA)^16^, elongation factor-G (EF-G), ternary complex (EF-Tu-aminoacyl-tRNA) and EF-Ts (Fig. 1b). Consistent with previous reports, immediately following the addition of all components (within 3 s for 50 % of the molecules at 50 nM Cy5-50S) subunit joining was observed by the appearance of a Cy3-Cy5 FRET signal (Fig. 1c and Extended data Fig. 1b)^17-20^. In 55 % of the mRNA molecules, the 50S subunit binds in a non-rotated high-FRET state while for 45 % of the molecules, we observe a semi-rotated state during subunit joining in agreement with IF2 being still bound to the ribosome^21-23^. Translation elongation at the single amino-acid resolution can be monitored by the 30S-h44/50S-h101 FRET pair, which reports on the two main steps of translation elongation: peptidyl transfer and translocation (Fig. 1b), where the ribosome transitions from a non-rotated, high-FRET state to a rotated, low-FRET state, respectively^15,16^. We monitored translation of individual codons over several minutes and reliably quantified the lifetime in the rotated and non-rotated states at up to 500 nM aminoacyl-tRNA (aa-tRNA) and 160 nM EF-G concentrations (Fig 1d, e) with values in agreement with previous single-molecule experiments^18,24^.

Simultaneously to translation elongation, we monitored transcription progression by following the DNA template signal. Upon delivery of all four NTPs, transcription immediately resumes (Extended Data Fig. 9) and could be monitored over several minutes^14^. Transcription termination is characterized by a single-step disappearance of the Cy3.5 signal. Plotting the time from NTP addition till DNA template dissociation for individual molecules allowed us to calculate average transcription rates (Fig. 1f), which are NTP concentration dependent and in agreement with previous bulk in vitro transcription and single-molecule studies as well as in vivo rates^8,14,25,26^. Thus, our reconstitution of the combined transcription-translation system recapitulates the activities of the systems previously studied in isolation.

### Stalled RNAP leads to ribosome slowdown upon collision

The full control of all components in our single-molecule assays allows us now to compare the translation elongation kinetics of single ribosomes either translating unhindered on an mRNA (if the RNAP transcribes faster than the ribosome translates the mRNA) or in the presence of a paused RNAP leading to a collision between the ribosome and the RNAP. We used a DNA template that allowed us to stall the ribosome and the RNAP with 46 nucleotides (nt) apart between the ribosome P-site to the RNAP active site (Fig. 2a, Supplementary Table 1,2, methods). This state was previously structurally characterized and classified as a “coupled state TTC-B” (Fig. 2d)^11^. Based on the cryoEM structures, this coupled state allows the translation of maximally 6 amino acids until both machineries collide, a state that has been structurally characterized and classified as a “collided state” (TTC-A, 28 nt separating P-site to RNAP active site, Fig. 2d)^4,10,11^. Delivery of all translation elongation factors together with NTPs triggered simultaneous translocation of both machines (Fig. 2b). The actively transcribed mRNA was immediately getting translated and we were able to track translation of up to 31 codons, until the stop codon was reached (Fig. 2e).

**Fig. 2:**
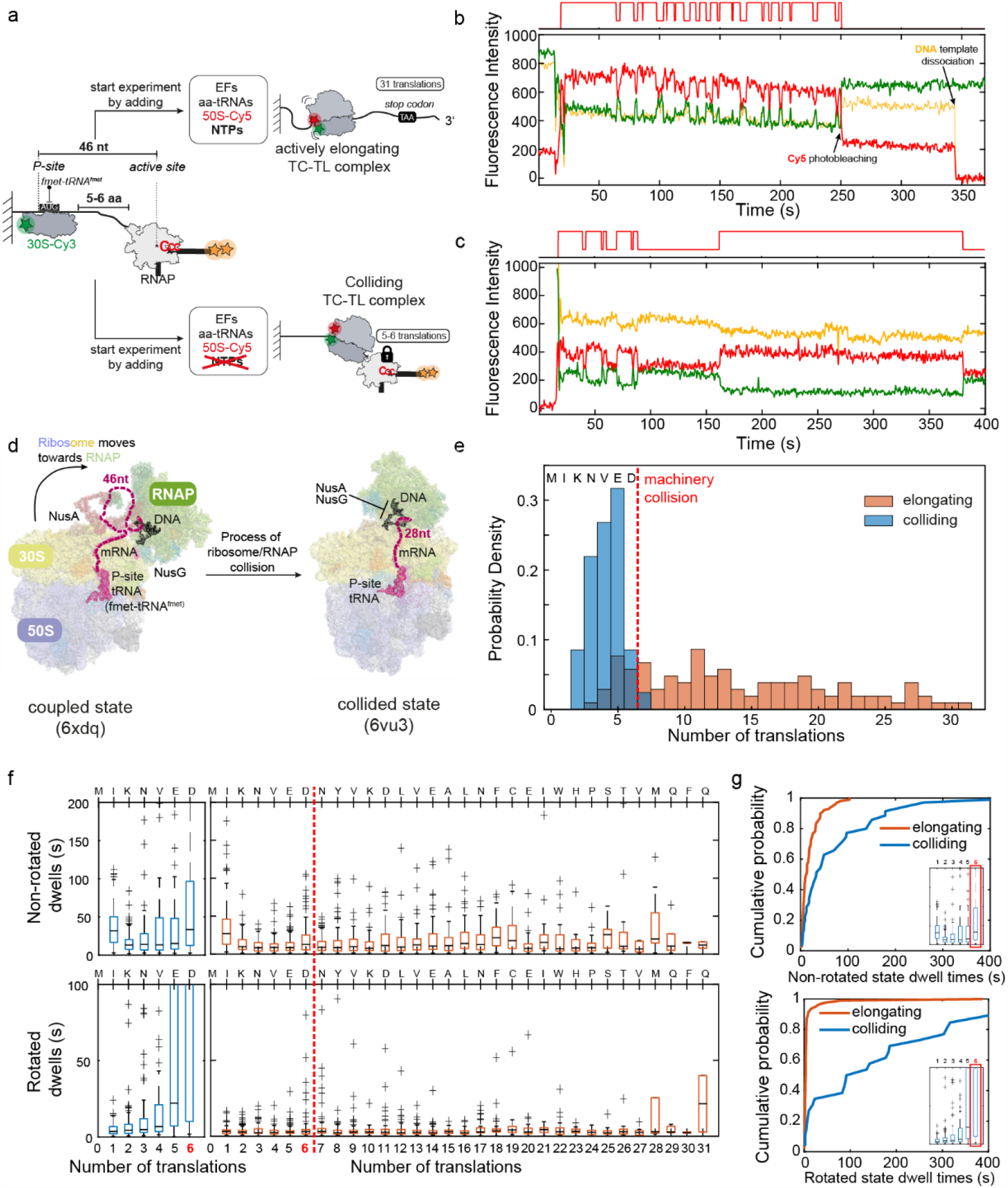
Ribosome slows down while colliding with RNAP. **a**, Experimental design. **b, c**, Representative traces for elongating (**b**) and colliding conditions (**c**). **d**, Expressome structures visualizing the start and end states of the ribosome slow-down upon colliding with the RNAP, represented by the coupled state (pdb: 6xdq) transitioning into the collided state (pdb: 6vu3). **e**, Probability density histograms for number of translations in colliding (blue, n=81) and elongating (orange, n=104) conditions. **f**, Boxplots of non-rotated (top) and rotated (bottom) dwell times during colliding (left, blue) and elongating (right, orange) conditions at 150 nM total aa-tRNAs and 50 nM EF-G. Note that at the given y-axis scale not all outliers are visible. **g**, Cumulative probability distribution of the dwell times for the 6^th^ amino acid incorporation for non-rotated state (top) and rotated state (bottom).

To investigate the effect of a stalled RNAP on ribosome translocation, we next repeated the experiment without delivery of NTPs (Fig. 2c). As expected, withholding NTPs during the single-molecule experiment led to collision of the ribosome into the artificially paused RNAP. Under these conditions, the largest fraction of ribosomes (32 %) completed 5 translation cycles and 9 % of the ribosomes completed 6 translation cycles before the end of the experiment or before photobleaching occurred (Fig. 2e). Comparing the median dwell times for translation of the individual codons of unhindered ribosomes with ribosomes that are in the process of colliding with the RNAP, showed that the ribosome slows down upon collision with the RNAP (Fig. 2f, compare blue versus orange data). The non-rotated dwell times gradually increase once the ribosome is four amino acids away from the RNAP (Fig. 2f and Extended Data Fig. 2c). This behavior is even more pronounced for the rotated state dwell times, which start to increase already at an RNAP to ribosome separation of 6 amino acids. In absence of a stalled RNAP, the dwell times for the non-rotated state (rate-limited by aa-tRNA arrival) or rotated-state (rate-limited by EF-G arrival) can be fitted to a single-exponential function in agreement with a single rate limiting step (Extended Data Fig. 2a)^27^. In contrast, under colliding conditions, the dwell times do not follow single-exponential kinetic behavior anymore suggesting additional rate-limiting steps while transitioning (Fig. 2d) from coupled to collided state (Fig 2g, Extended Data Fig. 2b-c). Lastly, we observe that under colliding conditions (Extended Data Fig. 1c and e), the ribosome preferentially ends up in the rotated state (63 % of the molecules compared to 24 % for elongating conditions), similarly to what has been found in vivo for a pseudouridimycin stalled expressome^28^.

### Extent of coupling depends on the length of the intervening mRNA

Available cryoEM structures represent either collided states or coupled states up to an intervening mRNA of 47 nt between the ribosome P-site and RNAP active site (6 amino acids inter-translation distance; Fig. 2d, left structure). Structures with longer intervening mRNA could so far not be obtained in vitro^10,11,13^. Furthermore, cryoET structures obtained from active expressomes in vivo did not show any density for the intervening mRNA^12^. This raises the question whether in vivo the ribosome and the RNAP can also physically couple if their intervening mRNA length is longer than visualized by current methods and what the kinetics of such coupling/uncoupling interactions would be.

To this end, we developed single-molecule assays to directly track in real-time the coupling and uncoupling dynamics between a stalled RNAP and a stalled ribosome at various lengths of intervening mRNA between both machineries (46-457 nt between P-site and RNAP active site; Fig. 3a). We labeled the ribosome and the RNAP with a donor and acceptor dye, such that FRET is only observed if both machineries are in a coupled state (Fig. 3b and Extended Data Fig. 3a). The ribosome was labeled at its mRNA-entry channel (rRNA extension at h33a^15,29^) with a donor Cy3-dye and the RNAP at the N-terminus or C-terminus of the β’-subunit with an acceptor Cy5-dye using ybbR-tag mediated dye-coupling (methods)^30,31^. Labeling both the ribosome^15,16,24^ and the RNAP (Extended Data Fig. 3b) at these positions did not detectably affect activity, and results in an inter-dye distance of 30-57 Å if the RNAP and the ribosome are in a coupled state (Extended Data Fig. 3a).

**Fig. 3:**
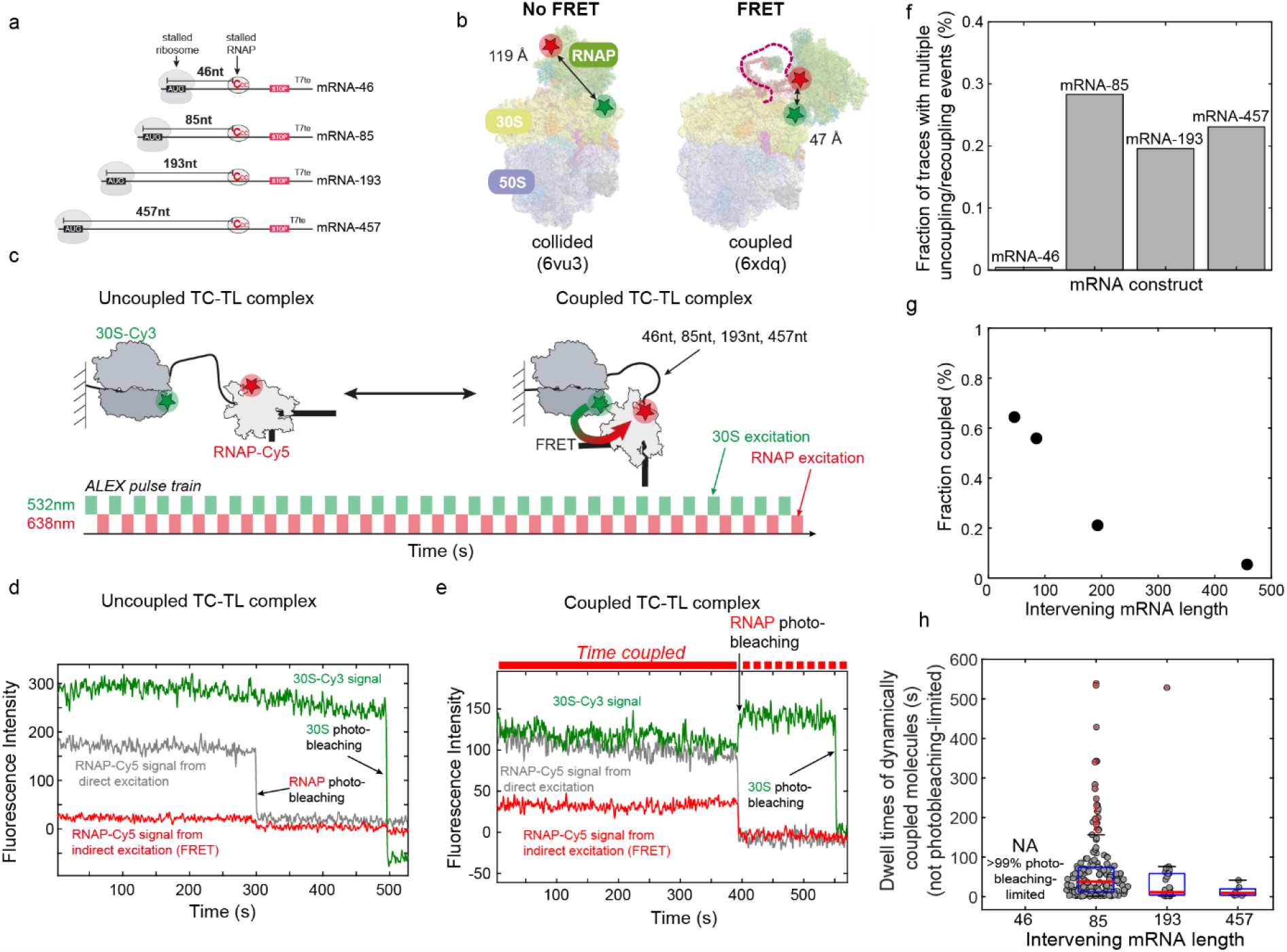
Real-time tracking of ribosome/RNAP coupling dynamics in dependence of intervening mRNA length. **a**, DNA template design. **b**, Location of labeling sites within expressome shown for collided state (left; pdb: 6vu3) and coupled state (right; pdb: 6xdq). **c**, Experimental smFRET setup for detecting ribosome/RNAP coupling kinetics in absence of translation and transcription elongation. **d, e**, Representative traces for uncoupled (left) and coupled (right) expressomes. **f**, Fraction of traces having multiple coupling/recoupling events as function of increasing mRNA length. **g**, Effect of intervening mRNA length on coupling efficiency. Number of evaluated molecules in **f** and **g**: n=340 (mRNA-46); n=297 (mRNA-85); n=218 (mRNA-193); n=239 (mRNA-457). **h**, Effect of intervening mRNA length on coupled lifetime of dynamically coupling molecules. Number of evaluated dwell times are: n=92 (mRNA-85); n=26 (mRNA-193); n=7 (mRNA-457).

After immobilizing the pre-assembled expressomes on a microscope slide (see methods), we used <u>a</u>lternating-laser <u>ex</u>citation (ALEX)^32^ at a wavelength of 532 and 638 nm (Fig. 3c-e), in order to distinguish uncoupling events from photobleaching (methods). For the shortest intervening mRNA (46 nt), we observe that 64 % of molecules are coupled and remain coupled till photobleaching (Fig. 3g), with a lower bound of 104 ± 2 s for the coupled lifetime (Extended Data Fig. 4b). This is in agreement with the formation of stable complexes as suggested from the cryoEM structures^10,11^. When increasing the intervening mRNA lengths between both machineries, we observed a gradual decrease in the fraction of molecules that are coupled at least once during the experiment, dropping to 5 % for 457 nt intervening mRNA (Fig. 3g and Extended Data Figure 5). We also find that 20-30 % of those coupled molecules show uncoupling/coupling dynamics during the experimental time (Fig. 3f, Extended Data Figure 4a and Extended Data Figure 5) with coupled dwell times decreasing with increasing intervening mRNA length (Fig. 3h). In addition, re-coupling becomes slower with increasing intervening mRNA lengths, taking in average 11.1 ± 0.4 s and 28 ± 3 s to recouple for the 85 nt and 193 nt intervening mRNA constructs (Extended Data Fig. 4c). Overall, these experiments demonstrate that ribosome and RNAP can physically interact even if the ribosome does not closely trail behind the RNAP but that the physical coupling efficiency decreases with increasing intervening mRNA length.

**Fig. 4:**
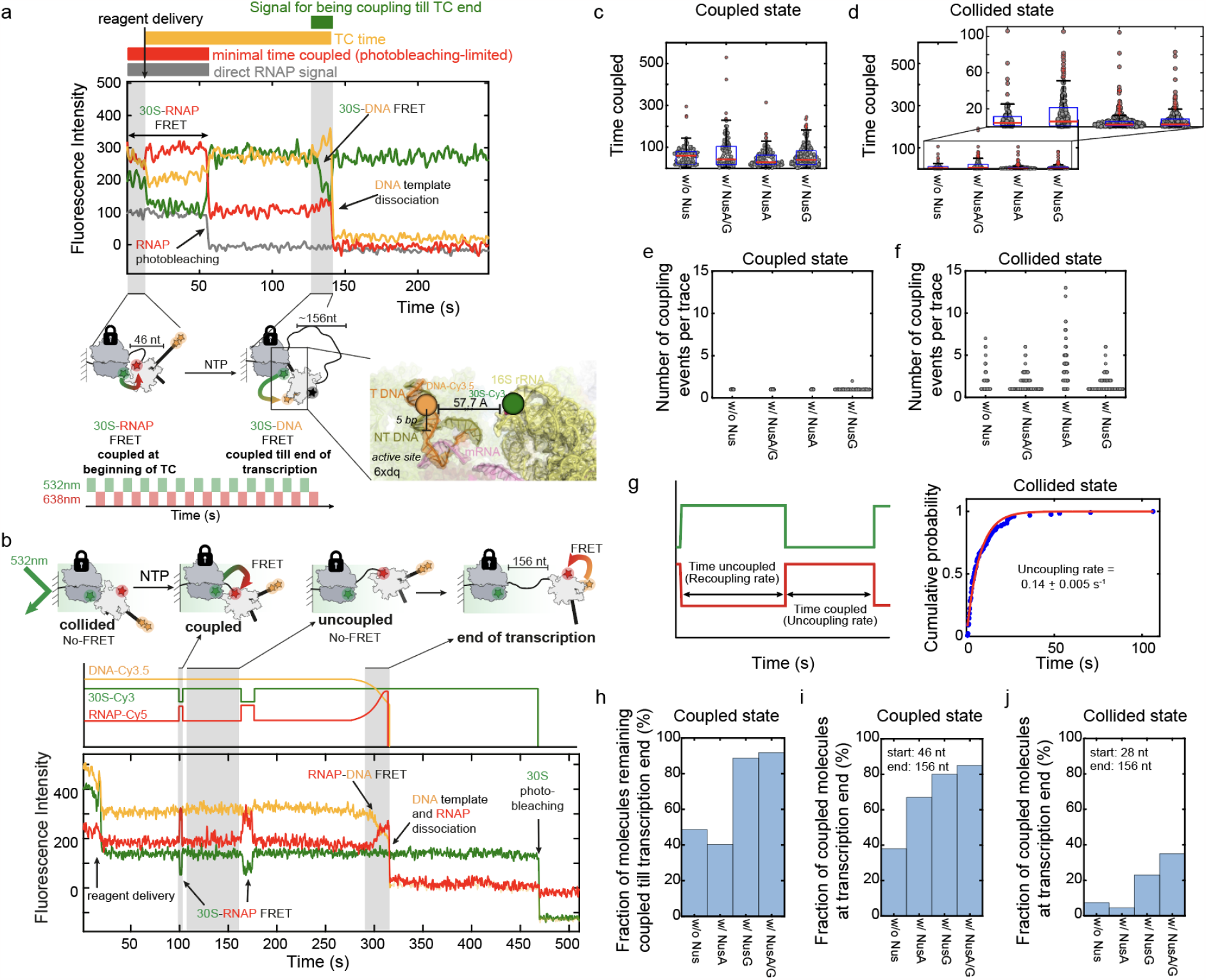
Nus factors increase coupling during active transcription elongation and RNAP/ribosome collisions reduce subsequent coupling efficiency. **a**, Experimental setup, example trace to track coupling during active transcription elongation and expressome structure displaying the distance of the 30S-Cy3 label in h33a (green) from the DNA-Cy3.5 (yellow-orange) label on the non-template DNA strand (pdb:6xdq). See main text for more details. **b**, Representative trace for transcription starting from collided state. Idealized trace is shown on top. Expressome species occurring during the experiment are indicated. **c, d**, Beeswarm plots representing all coupled dwell times for all molecules during transcription from coupled (**c**) and from collided state (**d**). **e, f**, Number of coupling/recoupling events per trace during transcription from coupled (**e**) and collided state (**f**). Number of evaluated molecules in **c-f**: n=204 (without factors); n=224 (with NusA/G); n=215 (with NusA); n=197 (with NusG) for transcription out of coupled state and n=75 (without factors); n=162 (with NusA/G); n=107 (with NusA); n=161 (with NusG) for transcription out of collided state. **g**, Coupled dwell times during transcription following collision fitted to 1-exponential function. Evaluated signal is shown on the left. **h**, Fraction of molecules remaining coupled to the end of transcription. **i, j**, Fraction of molecules coupled at the end of transcription starting from coupled (**i**) or collided state (**j**). Number evaluated molecules in **h** and **i** is identical to **c** and **e**. Number of evaluated molecules in **j**: n=174 (without factors); n=140 (with NusA/G); n=110 (with NusA); n=196 (with NusG).

**Fig. 5:**
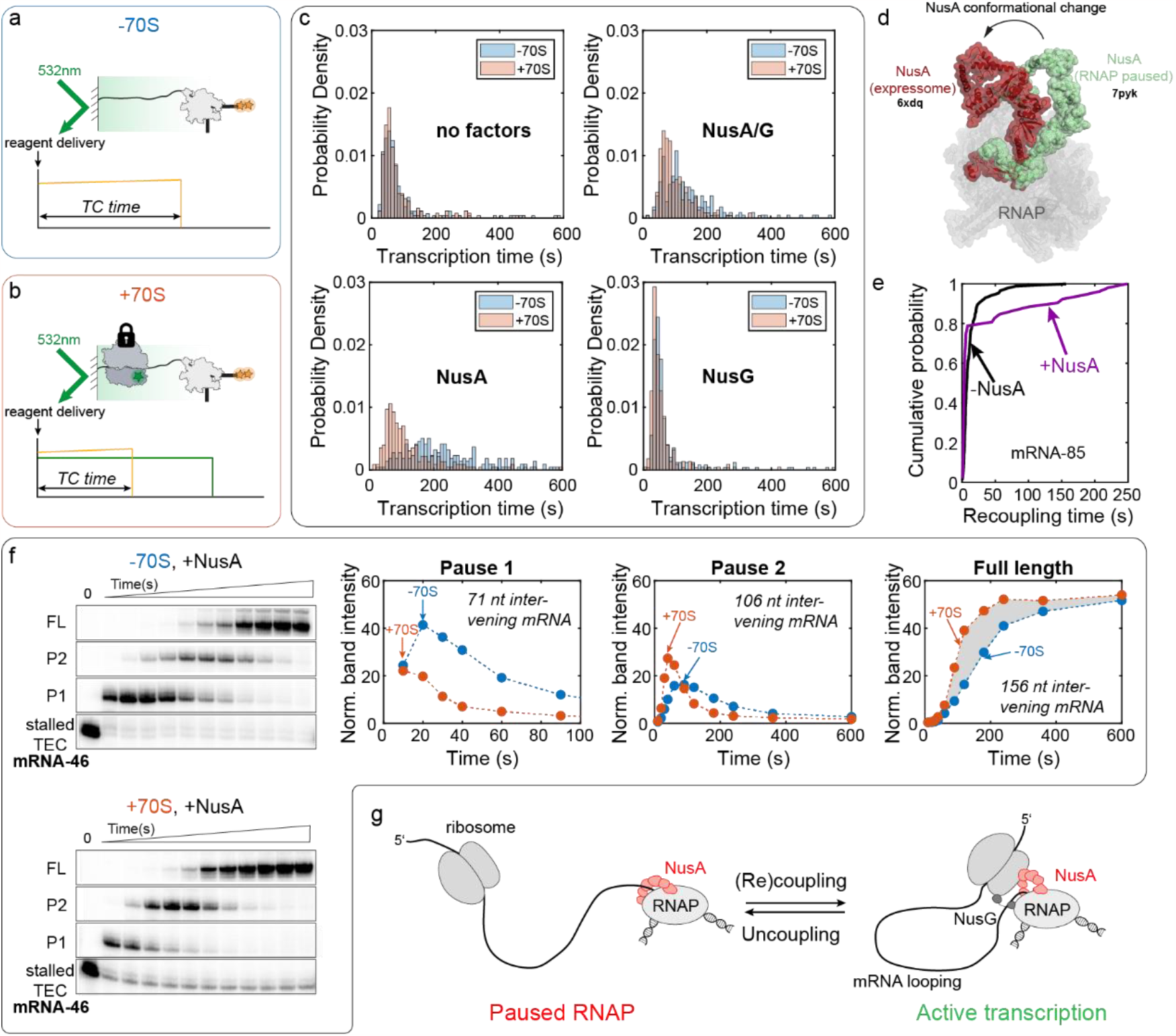
Ribosome reactivates NusA-paused RNAP via mRNA looping. **a, b**, Overview of single-molecule experimental setup. Start of experiment was triggered by delivery of 50 μM NTPs and Nus factors (no factors, with NusA/NusG, with NusA, with NusG; at 1 μM each) to either immobilized stalled mRNA-46 TEC (−70S, (**a)**) or stalled expressome (+70S, **(b)**). **c**, Overlay of probability density distribution for single-molecule transcription times without 70S (blue) and with 70S (orange) in presence and absence of Nus factors. Number of evaluated molecules (−70S/+70S): n=196/208 (without factors); n=223/243 (with NusA/G); n=218/227 (with NusA); n=254/222 (with NusG). **d**, NusA conformations in paused RNAP-bound (pdb: 7pyk) and in expressome-bound (pdb: 6xdq) cryoEM structures. The structures were aligned on the RNAP C-alpha backbone atoms using PyMol. **e**, Cumulative probability density for NusA-dependent or -independent expressome recoupling lifetimes observed for the mRNA-85 expressome in absence of transcription. **f**, Left: Single-round transcription assays with mRNA-46 in presence of NusA and with or without 70S analyzed by denaturing gel electrophoresis. Stalled TEC, pause 1, pause 2 and full-length mRNA bands are shown. Top right: Normalized band intensities are plotted as a function of time and shown for pause 1, pause 2 and full-length mRNA. **g**, Model for ribosome-assisted RNAP-activation via long-range interactions.

### Nus factors drive coupling during active transcription

Next, we investigated the effect of transcription elongation on the coupling efficiency. To this end, we immobilized expressomes with 46 nt intervening mRNA sequence and containing Cy5-RNAP, Cy3-30S and the 3’-end of the DNA template labeled with 2xCy3.5. Under these conditions, 78 % of the molecules showed a Cy3-Cy5 FRET signal at the start of the experiment, characteristic for the presence of coupled complexes (Fig. 4a, Extended Data Fig. 3a, Extended Data Fig. 6a and Extended Data Fig. 7). After starting single-molecule imaging, we added NTPs to initiate transcription elongation, but kept the ribosome stalled at the RBS by not adding translation elongation factors. 37 % of the traces showed a 30S-Cy3 to RNAP-Cy5 FRET signal till the end of transcription without any uncoupling or recoupling events (Fig. 4e and Extended Data Fig. 7a) demonstrating constant coupling between ribosome and RNAP during active transcription elongation. However, for the remaining 63 % of the molecules, the RNAP-Cy5 signal photobleached before the end of transcription (Fig. 4a and Extended Data Fig. 7b), allowing us to only provide a lower estimate of 59.3 s for the median coupled lifetime during active transcription elongation (Fig 4c). Interestingly, for a fraction of the traces (38%; Fig 4i), we observe the appearance of FRET signal between the 30S-Cy3 and the DNA-Cy3.5 at the end of the 156-nucleotide mRNA transcription (Fig. 4a). This FRET, for which both dyes are not affected by photobleaching under our experimental conditions, shows that the ribosome is positioned close to the end of the DNA template, a configuration that can only be achieved when the two machines are coupled at the end of transcription (Fig. 4a). Using this Cy3/Cy3.5 FRET signal as a proxy for coupling once the full-length mRNA has been transcribed (156 nucleotides intervening mRNA sequence), we next compared the effect of the different Nus-factors on coupling efficiency. We find that in presence of NusG (+/- NusA), the fraction of molecules that remain coupled to the end of transcription is approximately 2-fold higher, reaching over 90 % in presence of both factors (Fig 4h). Overall, our data shows that RNAP and ribosome can remain coupled during active transcription elongation, supported by transcription factor NusG, and that hundreds of nucleotides of mRNA can thereby be looping-out between both machineries.

### Coupling following ribosome/RNAP collisions becomes less efficient

Next, we wondered whether ribosome/RNAP collisions may influence the subsequent coupling efficiency. To this end, we immobilized a collided expressome (RNAP-Cy5, 30S-Cy3) to the imaging surface. In contrast to the coupled expressome, the collided expressome shows no FRET between RNAP-Cy5 and 30S-h33a-Cy3 (>100 Å between both dyes; Extended Data Fig. 3a and Fig 4b). We triggered transcription elongation of the RNAP by delivering NTPs (Fig 4b, Extended Data Fig. 6b). Transcription from the collided state immediately resumes after addition of all four NTPs (stalled TEC band completely chased in <10 s; see Extended Data Fig. 8), in agreement with previous studies^4^. As transcription proceeds and the RNAP transitions from collision to coupling with the ribosome, this transition shows as an appearance of a Cy3-Cy5 FRET signal (Fig 4b). Remarkably, coupling following a collision is >10 fold shorter compared to coupling in absence of a prior collision (Fig. 4c,d). The coupled lifetime reduces to only 7.0 ± 0.3 s (Fig. 4g) with the RNAP and the ribosome dynamically coupling and uncoupling multiple times during transcription elongation (Fig. 4f and Extended Fig. 6b). Interestingly, during some of these coupled states, we see dynamic FRET changes, suggesting transitions between multiple different coupled sub-states as was also seen in recent cryoEM structures^11^ (Fig. 4b). The fraction of traces that show coupled ribosome/RNAP at the end of transcription reduces to 7.4 % (versus 38 % without prior ribosome/RNAP collision; compare Fig 4j and Fig 4i) and increases to 23/35% in presence of NusG or both Nus-factors, respectively (compared to 80/85% without prior ribosome/RNAP collision). Overall, this demonstrates that RNAP/ribosome collisions reduce subsequent coupling efficiency. We hypothesize that the transient nature of coupling following a ribosome-RNAP collision may be attributed to a structural or compositional change of the ribosome/RNAP interface upon collision, such as loss of the omega subunit as suggested by the missing electron density for this RNAP subunit in the cryoEM maps of the collided state^10,11^.

### Coupled ribosome rescues NusA-paused RNAPs

Having demonstrated that the ribosome and the RNAP can physically couple also with long intervening mRNA, this raises the question whether this long-range physical coupling is also functional, in other words, can the ribosome activate a stalled RNAP also by long-range coupling interactions.

Therefore, we immobilized stalled transcription elongation complexes either in absence of a ribosome or in presence of ribosomes in a coupled state (mRNA-46). We started the single-molecule imaging reactions by the addition of NTPs and measured the time it takes to transcribe full-length mRNA molecules (Fig. 5a-b). In line with previous bulk experiments^33-35^, our single-molecule data (Fig. 5c) and bulk in vitro transcription reactions (Extended Data Fig. 9c,d) show a significant effect of NusA and NusG on transcription pausing or transcription activation, respectively, demonstrating the activity of the Nus-factors on transcription elongation. In absence of Nus-factors (median transcription time: 66.6 s (−70S) versus 59.8 s (+70S); 1.1-fold increase) or with NusG alone (49.2 (−70S) versus 42.7 s (+70S); 1.2-fold increase), we only see marginal ribosome-induced acceleration of transcription speed. With both Nus factors (116.0 s (−70S) versus 85.4 s (+70S); 1.4-fold increase) we observe a modest increase in median transcription time. However, for NusA alone (205.9 s (−70S) versus 93.3 s (+70S); 2.2-fold increase), transcription is significantly faster in presence of the ribosome, indicating that a coupled ribosome can rescue a NusA-paused RNAP.

In order to verify our single-molecule experiments, we performed bulk in vitro transcription assays. For this purpose, we purified the expressome (formed with mRNA-46) using streptavidin beads by immobilizing via biotin-labeled 50S subunits. This allowed us to enrich fully assembled expressomes in our transcription assays (see methods). While we again see only a slight increase in transcription speed for most conditions (without factors, NusG, NusA/G) in presence of the ribosome (Extended Data Fig. 9a-d, g), we see again the most significant ribosome-induced increase in transcription efficiency in presence of NusA alone (Fig. 5f and Extended Data Fig. 9a-e, g). We find two prominent pause sites for mRNA-46 that most probably are induced by formation of pause hairpins (Extended Data Fig. 9f)^36^. Importantly, these transcription assays allow us to determine pause-escape lifetimes in presence and absence of the ribosome and at which intervening mRNA distance between ribosome and RNAP the ribosome-accelerated pause release occurs (methods). We find that at the first pause site (at 134 nt, 71 nt intervening mRNA), the ribosome decreases the pause escape lifetime 1.35-fold from 69 ± 2 s to 51 ± 4 s in presence of NusA (Extended Data Fig. 9e). In case of the second pause site (at 169 nt, 106 nt intervening mRNA) we observe a stronger effect (ribosome decreases the pause escape lifetime 2.6-fold from 188 ± 5 s to 72 ± 2 s).

Structurally, both NusA-mediated RNAP pausing^33^ as well as NusA-associated coupling in the expressome were described^11^. RNAP-bound NusA needs to be re-positioned significantly prior to docking to the ribosome, as the NusA conformation significantly differs between expressome and NusA-bound RNAP structures (Fig. 5d). Remarkably, our single-molecule data capture the slow kinetics of NusA repositioning upon re-coupling. While the recoupling rates in absence of Nus factors follow mostly single exponential behavior (96 % fast on-rate of 0.097 ± 0.03 s^-1^), recoupling in presence of NusA alone is more complex (Fig. 5e, Extended Data Fig. 10). Here, the data consistently deviate from pure single-exponential behavior and we observe additional, slower components describing the NusA-dependent re-coupling rates (Fig. 5e, Extended Data Fig. 10). Overall, these data are consistent with a model in which long-range coupling between the ribosome and a NusA-mediated paused RNAP re-positions NusA on the RNAP from a pause-promoting state to an activated state.

## Discussion

Transcription-translation coupling has served as a paradigm on how macromolecular machines cooperate inside cells. By in vitro-reconstituting the complete active transcription-translation machinery and using multi-color single-molecule imaging, we have simultaneously tracked transcription and translation elongation, together with the physical and functional coupling between both machineries. Decades of work and recent cryoEM/ET structures of the expressome suggest that transcription-translation coupling is general^1,8-13,37,38^, but recent data also suggest that coupling is more stochastic in *E. Coli*^39,40^ or that the two processes are mostly decoupled^41^. Our work consolidates both models as we show that physical and functional coupling can also occur efficiently when both machines are far apart along the mRNA sequence, by looping of the mRNA. Our new model has several major implications for the field and beyond.

Ribosome-RNAP collisions have been suggested to be a possible mechanism in vitro for the ribosome to rescue a back-tracked RNAP from pausing^4,5^. However, no collided ribosome/RNAP complexes were detected in vivo in absence of antibiotics^12^, which can be explained by our findings that the ribosome slows down before colliding into the RNAP, making ribosome/RNAP collisions less likely to occur frequently. Instead, more efficient ribosome-induced RNAP activation via long-range coupling can stochastically occur during the entire transcription process, over several hundred nucleotides separation, therefore relevant for an average *E. Coli* mRNA length of 1000 nucleotides^42^. Long-range physical coupling may also occur between the RNAP and a non-leading ribosome. Such a mechanism would provide a means for the RNAP to sense the ribosome loading occupancy on the mRNA rather than just the physical proximity of the leading ribosome, because a higher number of ribosomes on the nascent mRNA would provide more chances for long-range RNAP/ribosome encounters and thereby increase RNAP activation.

We show that the mRNA can loop-out for several hundred nucleotides and that coupling remains stable during active transcription, further supported by NusG and NusA. This requires rethinking our understanding of how the ribosome modulates Rho-dependent transcription termination^43,44^. In a model, where the mRNA can loop-out by more than 100 nucleotides, Rho could also engage with the mRNA (Rho/mRNA foot-print of 57/85 nucleotides^45^) while the RNAP and the ribosome are physically coupled. Whether Rho could push away the ribosome in order to reach the RNAP for terminating transcription and how this is coordinated with NusG remains to be investigated.

Co-transcriptional RNA looping is likely a more general feature for increasing the efficiency of transcription-coupled processes^46^. Both in bacterial rRNA transcription (mediated by interactions between the ribosomal RNA transcription antitermination complex and the RNAP^47^) and eukaryotic co-transcriptional splicing (mediated by a U1 snRNP – Pol-II interactions^48^) transcription-mediated RNA looping could provide means for increased RNA processing efficiency (RNase III processing and splicing, respectively).

Overall, our work underlines the importance of studying biological processes not in isolation but in context of each other and highlights the power of multi-color single-molecule experiments to image complex active reconstituted systems for maximal control of all the components and for tracking the dynamics of complex multi-step processes at highest spatial and temporal resolution.

## Methods

### Strains and plasmids

Ribosome mutant SQ380 strains (B68 strain=30S helical extension in h33a, C68 strain=30S helical extension in h44 and ZS22 strain=50S helical extension in h101) and all plasmids for initiation and elongation factors were a kind gift from the Puglisi lab^15,16^. The plasmid for overexpressing initiator tRNA^fmet^ (pBStRNAfmetY2) was a kind gift from Emmanuelle Schmitt^49^. The plasmids for *E. Coli* RNAP (pVS10) and σ70 (pIA586) were purchased from Addgene (#104398 and #104399, respectively)^50^. The ybbR-peptide tag (DSLEFIASKLA)^51,52^ was cloned into the pVS10 plasmid by introducing the ybbR DNA sequence with primers, followed by 5’-end phosphorylation with T4 PNK (New England Biolabs, NEB) and simultaneous blunt ligation of the plasmid using T4 DNA ligase (NEB). For the N-terminal ybbR-mutant the tag was inserted between E15 and E16 in the β’-subunit; for the C-terminal β’-mutant it was inserted between E1377 and G1402, thereby deleting the region A1378-L1401. Gene sequences of ribosomal S1 protein, NusA, NusG, methionyl-tRNAfmet formyltransferase (MTF) and methionine tRNA synthetase (MetRS, truncated at K548 to obtain the monomeric form of the enzyme^53^ were cloned into a pESUMO vector backbone using Gibson assembly. Gene sequences were obtained from ASKA collection plasmids^54^.

### Sample preparation

*E. Coli* ribosomal subunits, initiation factor IF2 and elongation factors (EF-Tu, EF-G and EF-Ts) were prepared and purified as previously described^15,16^. WT RNAP and mutant versions were purified as described before^50^. Initiator tRNA^fmet^ was prepared and purified following the protocol by Mechulam et al.^55^. r-protein S1, NusA and NusG were overexpressed as N-terminal fusions with His_6_-SUMO. They were purified by lysing the cells in IMAC buffer (50 mM TrisHCl pH 8.0, 300 mM NaCl, 10-20 mM imidazole, 10 mM 2-mercaptomethanol) using a microfluidizer, loading the clarified lysate on 5 mL HisTrap HP columns (Cytiva) and eluting with increasing imidazole concentrations over 20 column volumes. Protein fractions were pooled and the fusion tag was cleaved overnight with His-tagged Ulp1 protease, while dialyzing against IMAC buffer without imidazole. The cleaved tag was removed by a second HisTrap purification. Protein fractions were pooled, concentrated using 15 mL Amicon Ultracel 10 or 30K concentrators and further purified via size exclusion chromatography using either HiLoad S75 (NusG) or HiLoad S200 (NusA, S1) columns (Cytiva) equilibrated with storage buffer (S1 in 50 mM HEPES-KOH pH 7.6, 100 mM KCl, 1 mM DTT and NusA/NusG in 20 mM TrisHCl pH 7.6, 100 mM NaCl, 0.5 mM EDTA/DTT). Protein fractions were pooled and concentrated using 15 mL Amicon Ultracel 10 or 30K concentrators to ∼100 μM. Aliquots were flash-frozen with liquid nitrogen and stored at -80 °C. In case of MTF and MetRS, the preparation was done as described above with following changes (and adapted from Mechulam et al. and Schmitt et al.)^55,56^: (i) MTF IMAC buffer contained 10 mM K_2_HPO_4_/KH_2_PO_4_ pH 7.3, 100 mM KCl, 20 mM imidazole and 10 mM 2-mercaptoethanol; MetRS buffer contained 10 mM K_2_HPO_4_/KH_2_PO_4_ pH 6.7, 50 mM KCl, 20 mM imidazole, 100 μM ZnCl_2_ and 10 mM 2-mercaptoethanol. (ii) after a 2^nd^ HisTrap purification, MetRS containing fractions were pooled and dialyzed against storage buffer (10 mM K_2_HPO_4_/KH_2_PO_4_ pH 6.7, 10 mM 2-mercaptoethanol and 50 % (v/v) glycerol). Aliquots were flash-frozen with liquid nitrogen and stored at -80 °C. (iii) after a 2^nd^ HisTrap purification, MTF was further purified via a 5 mL HiTrap Q FF column (Cytiva) and eluted using a linear gradient of 20 - 100 % into Q-sepharose buffer (10 mM K_2_HPO_4_/KH_2_PO_4_ pH 7.3, 500 mM KCl and 10 mM 2-mercaptoethanol). Protein fractions were pooled and dialyzed against storage buffer (10 mM K_2_HPO_4_/KH_2_PO_4_ pH 6.7, 100 mM KCl, 10 mM 2-mercaptoethanol and 50 % (v/v) glycerol). Aliquots were flash-frozen with liquid nitrogen and stored at -80 °C.

### Charging tRNA^fmet^ and elongator tRNAs

Typically, initiator tRNA^fmet^ (20 μM) was simultaneously charged and formylated in 800 μL reactions using 100 μM methionine, 300 μM 10-formyltetrahydrofolate (10-TFHFA), 200 nM MetRS and 500 nM MTF in charging buffer (50 mM TrisHCl pH 7.5, 150 mM KCl, 7 mM MgCl_2_, 0.1 mM EDTA, 2.5 mM ATP and 1 mM DTT), incubating for 5 min at 37 °C^55,57^. The fmet-tRNA^fmet^ was immediately purified by addition of 0.1 volume sodium acetate (NaOAc, pH 5.2), extraction with aqueous phenol (pH ∼ 4) and precipitation with 3 volumes of ethanol. The tRNA pellet was solubilized in ice-cold tRNA storage buffer (10 mM NaOAc pH 5.2, 0.5 mM MgCl_2_) and further purified with Nap-5 column (Cytiva) equilibrated with the same buffer. The eluate was aliquoted, flash-frozen with liquid nitrogen and stored at -80 °C. Elongator tRNAs were purchased (tRNA MRE600, Roche) and charged typically at 500 μM concentration in presence of 0.2 mM amino acid mix (each), 10 mM phosphoenolpyruvate (PEP), 20 % (v/v) S150 extract (prepared following published protocol)^58^, 0.05 mg/mL pyruvate kinase (Roche) and 0.2 U/μL thermostable inorganic pyrophosphatase (TIPP, NEB) in total tRNA charging buffer (50 mM TrisHCl pH 7.5, 50 mM KCl, 10 mM MgCl_2_, 2 mM ATP and 3 mM 2-mercaptoethanol)^29^. Typically, 200 μL reactions were incubated at 37 °C for 15 min and then immediately purified as described above. To remove NTP contaminations introduced by the S150 extract, the aa-tRNAs were further purified over a S200 increase column (Cytiva), pre-equilibrated in tRNA storage buffer. The eluted fractions were combined, concentrated with 2 mL Amicon Ultracel 3K concentrators, aliquoted and flash-frozen with liquid nitrogen and stored at -80 °C. Charging efficiency (typically >90 %) was verified with acidic urea polyacrylamide gel electrophoresis (PAGE) as described by Walker et al.^57^.

#### Dye-labeling of expressome components

Hairpin loop extensions of mutant ribosomal subunits were labeled with prNQ087-Cy3 (30S) or prNQ088-Cy5 (50S) DNA oligonucleotides, complementary to mutant helical extensions^15,16,18^. Just prior to the experiments, each subunit was labeled separately at 2 μM concentration using 1.2 equivalents of the respective DNA oligonucleotide by incubation at 37 °C for 10 min and then at 30 °C for 20 min in a Tris-based polymix buffer (50 mM Tris-acetate pH 7.5, 100 mM potassium chloride, 5 mM ammonium acetate, 0.5 mM calcium acetate, 5 mM magnesium acetate, 0.5 mM EDTA, 5 mM putrescine-HCl and 1 mM spermidine). RNAP-ybbR mutants (coreenzyme) were labeled by mixing 7 μM RNAP, 14 μM SFP synthetase and 28 μM CoA-Cy5 dye in a buffer containing 50 mM HEPES-KOH pH 7.5, 50 mM NaCl, 10 mM MgCl_2_, 2 mM DTT and 10 % (v/v) glycerol^59^. Typically, 100 μL reactions were incubated at 25 °C or 37 °C for 2 h and analyzed on denaturing protein gels. The holoenzyme was formed by incubating 1.11 μM Cy5-labeled RNAP with 3 equivalents σ70 for 30 min on ice in RNAP storage buffer (20 mM TrisHCl pH 7.5, 100 mM NaCl, 0.1 mM EDTA, 1 mM DTT, 50 % (v/v) glycerol). Aliquots were stored at -20 °C. SFP and free dye were removed on imaging surface prior to experiments.

DNA templates were purchased (TwistBioscience) and amplified via PCR using p0030 forward and p0075 reverse abasic primers, generating single-stranded 5’-overhangs for both DNA strands^14^. The fragments were purified on 2 % agarose gels, extracted using a QIAGEN gel extraction kit and buffer exchanged in e55 buffer (10 mM Tris-HCl pH 7.5 and 20 mM KCl) using 0.5 mL Amicon Ultracel 30K concentrators. The 5’-overhang of the template DNA was hybridized by mixing with 1.2 equivalents of p0088-2xCy3.5 DNA oligonucleotide at 68 °C for 5 min, followed by slow cool down (∼1 h) to room temperature.

#### Single-round in vitro transcription assays

Stalled transcription elongation complexes (TECs) were assembled in transcription buffer (50 mM Tris-HCl pH 8, 20 mM NaCl, 14 mM MgCl_2_, 0.04 mM EDTA, 40 μg/mL non-acylated BSA, 0.01 % (v/v) Triton-X-100 and 2 mM DTT) essentially as described previously^14,60^. In brief, 50 nM DNA template was incubated (20 min, 37 °C) with four equivalents of RNAP in presence of 100 μM ACU trinucleotide, 5 μM GTP, 5 μM ATP (+ 150 - 300 nM ^32^P α-ATP, Hartmann Analytic), halting the polymerase at U24, to prevent loading of multiple RNAPs on the same DNA template. Next, re-initiation of transcription was blocked by addition of 10 μg/mL rifampicin. The RNAP was walked to the desired stalling site by addition of 10 μM UTP and incubating at 37 °C for 20 min. The 30S ribosomal subunit was loaded for 10 min at 37 °C by incubating 25 nM stalled TEC with 250 nM 30S (B68 mutant, pre-incubated with stoichiometric amounts of S1 protein for 5 min at 37 °C) in presence of 2 μM IF2, 1 μM fmet-tRNA^fmet^ and 4 mM GTP in polymix buffer with 15 mM Mg(OAc)_2_. To enrich for fully assembled TC-TL complexes the expressome was purified via immobilizing the 50S subunit (ZS22) onto streptavidin magnetic beads (NEB). For this, the 50S subunit was pre-annealed on h101 with prNQ302-prNQ303-prNQ304-p0109-biotin DNA oligonucleotide. The prNQ303 and prNQ304 DNA oligonucleotides contain a BamHI cleavage site to elute the purified TC-TL complex from streptavidin beads. 50S loading onto the stalled TEC loaded with 30S PIC occurred simultaneously while immobilizing the TC-TL complex on streptavidin magnetic beads (pre-equilibrated with polymix buffer with 15 mM Mg(OAc)_2_) for 10 min at room temperature. Typically, 50 μL beads were loaded with a total volume of 150 μL TC-TL complex. The immobilized complex was washed once with polymix buffer with 15 mM Mg(OAc)_2_ and then eluted with 100 μL polymix buffer with 15 mM Mg(OAc)_2_, while cleaving with BamHI for 20 min at 37 °C. The eluate was chased with 50 μM NTPs in presence or absence of Nus factors (at 1 μM each, when present), 4 mM GTP, polymix buffer with 15 mM Mg(OAc)_2_ and 100 mM potassium glutamate (KGlu). Time points were taken before NTP addition (t=0 s) and at 10, 20, 30, 40, 60, 90, 120, 180, 240, 360 and 600 s for each condition, mixing 2.5 μL sample with 5 μL stop buffer (7 M urea, 2xTBE, 50 mM EDTA, 0.025 % (w/v) bromphenolblue and xylene blue) and incubated at 95 °C for 2 min. As size reference, the ssRNA ladder (NEB, sizes: 50, 80, 150, 300, 500 and 1000 nt) was 5’-end labeled with ^32^P γ-ATP. The reactions were analyzed on 6 % denaturing PAGE (7 M urea in 1xTBE), running in 1xTBE and 50 W for 2-3 h. Gels were dried, exposed overnight on a phosphor screen (Cytiva, BAS IP MS 2040 E) and imaged using a Typhoon FLA 9500. Bands (P) were integrated using ImageLab software (BioRad) and divided by the total RNA per lane (T) to compensate for pipetting errors, as described by Landick et al.^61^. Normalized band intensities (P/T) were plotted as function of time. Pause-escape lifetime were essentially fitted as described by Landick et al. by plotting ln(P/T) against time and fitting pause-escape data range (indicated in plots) with a linear equation (y=m*x+b, with m being the pause-escape rate).

#### Single-molecule transcription-translation-coupling assays

Stalled TECs were essentially formed as described for single-round transcription with following changes: (i) Typically, labeled DNA was used (pre-annealed with p0088-2xCy3.5). (ii) Stalled transcription was performed in presence of 10 μM ATP and 10 μM GTP. (iii) Re-initiation of transcription was blocked with 1 mg/mL heparin (final concentration). (iv) For immobilization, the 5’-end of the nascent mRNA was labeled with biotin-DNA oligonucleotide (prNQ127-p0109-biotin). (v) When necessary, Cy5-labeled RNAP was used (N-terminal ybbR-tag for prNQ215, prNQ216 and prNQ219; C-terminal ybbR-tag for prNQ219, prNQ291 and prNQ301). Typically, ribosomes (250 nM 30S, 500 nM 50S (when present)) were loaded for 5 min at 37 °C by incubating 4 nM stalled TEC in presence of 2 μM IF2, 1 μM fmet-tRNA^fmet^ and 4 mM GTP in polymix buffer with 15 mM Mg(OAc)_2_. The loading reaction was diluted to 200-800 pM stalled TEC concentration (DNA template-based) with an IF2-containing (2 μM) polymix buffer at 15 mM Mg(OAc)_2_ and immobilized on biotin-polyethylene-glycol (PEG) functionalized slides coated with NeutrAvidin for 10 min at room temperature^62^. Unbound components were washed away with an IF2-containing (2 μM) imaging buffer (polymix buffer with 15 mM Mg(OAc)_2_, 100 mM KGlu) and imaging was immediately started. The imaging buffer was supplemented with an oxygen scavenger system (OSC), containing 2.5 mM protocatechuic acid and 190 nM protocatechuate dioxygenase and a cocktail of triplet state quenchers (1 mM 4-citrobenzyl alcohol (NBA), 1 mM cyclooctatetraene (COT) and 1 mM trolox) to minimize fluorescence instability. For equilibrium experiments the imaging buffer in addition contained 4 mM GTP and 1 μM Nus factors (each, exact composition indicated in figures). For real-time experiments, the reaction was initiated while imaging (after 10 – 30 s), with delivery of 50 nM 50S-Cy5 (where applicable), 2 μM IF2, 10 – 160 nM EF-G (where applicable, in specified concentrations), 100 – 500 nM ternary complex (where applicable, in specified concentrations), 10 – 500 μM NTPs (each, where applicable, in specified concentrations) and 1 μM Nus factors (each, where applicable) in the same imaging buffer containing additional 4 mM GTP. The ternary complex was prepared as described previously^63,64^.

#### Instrumentation and analysis

We performed all single-molecule experiments at 21 °C using a custom-built (by Cairn Research: https://www.cairn-research.co.uk/), objective-based (CFI SR HP Apochromat TIRF 100XC Oil) total internal reflection fluorescence (TIRF) microscope, equipped with an iLAS system (Cairn Research) and Prime95B sCMOS cameras (Teledyne Photometrics). For standard TIRF experiments (3 colors), we used a diode-based (OBIS) 532 nm laser at 0.6 kW cm^-2^ intensity (based on output power). The fluorescence intensities of Cy3, Cy3.5 and Cy5 dyes were recorded at 5 frames per second (200 ms exposure time). For alternative laser excitation^32^ (ALEX) experiments, we operated the 532 nm laser at 0.73 kW cm^-2^ intensity and in every alternate frame illuminated the samples with a diode-based 638 nm laser (Omicron LuxX) at 0.12 kW cm^-2^ intensity (200 ms exposure time for each laser). Images were acquired using the MetaMorph software package (Molecular Devices) and single-molecule traces were extracted using the SPARTAN software package (v.3.7.0)^65^. Subsequent analysis was done with scripts^18,66^ written in MATLAB R2021a and previous versions (MathWorks). For data evaluation, only traces with single expressome molecules (containing 30S-Cy3 and DNA-2xCy3.5) were used. Transcription times were evaluated by assigning the time after reagent delivery till DNA template dissociation. Average transcription times (Fig. 1f) were obtained by fitting dwell time distributions to a convolution of a Gaussian (describes transcription) and exponential function (describes RNAP-stalling at the 3’-end before termination)^14^. For translation experiments, FRET transitions were assigned, selecting only for productive FRET states (High-FRET to Low-FRET state or vice versa). Translation counts were stopped when one of the dyes entered a dark state. The final translation events were photobleaching-limited. We kept those in for data evaluation, as they inform on ribosome stalling upon collision with the RNAP or upon encountering the stop codon. Lifetimes for rotated or non-rotated state were obtained by single-exponential fitting (in some cases (colliding conditions) double-exponential fitting) of cumulative dwell time distributions. For equilibrium coupling experiments, expressome molecules were first selected for assembled expressomes (30S-Cy3 signal, and direct excitation signal of RNAP-Cy5). Next, the coupling signal (30S-Cy3/RNAP-Cy5 FRET) was assigned. Cases, where no coupling occurred, went into the evaluation unassigned. Fraction of coupled molecules was calculated as: coupling efficiency = (number of coupled molecules with 30S-RNAP FRET)/(total number of expressome molecules). Uncoupling and recoupling rates were obtained by single- or double-exponential fitting of cumulative dwell time distributions. For real-time coupling experiments (ribosome stalled on RBS, RNAP chased with NTP) assembled expressome molecules were selected (30S-Cy3 signal, DNA-2xCy3.5 signal and in case of ALEX, direct excitation signal of RNAP-Cy5). When applicable, dwell times for transcription or coupling were extracted. For fraction of coupled molecules till end of transcription, we next assigned the characteristic signal (30S-Cy3/DNA-2xCy3.5 FRET) and calculated: fraction coupled = (number of coupled molecules with 30S-DNA FRET)/(total number of expressome molecules).

## Acknowledgements

We are grateful to Martyn Reynolds and Felix Jan Evers from Cairn Research, Marko Lampe from the EMBL Advanced Light Microscopy Facility, to Andrey Revyakin, members of the Puglisi lab, Thomas Hoffmann, Albert Tsai, Kavan Gor, Pratulya Shah and Eva Geissen for assistance in single-molecule data acquisition and processing. We thank Anastasiia Chaban for preparation of some NusA protein batches, David Will from the EMBL chemical core facility for initial assistance in slide preparation and the EMBL protein expression and purification core facility for purification of recombinant *E. Coli* RNAP. The authors thank Julia Mahamid for critical reading of the manuscript. We are grateful to Helga Grötsch, Jonas Weidenhausen, the Lemke lab and the entire Duss lab for assistance in the wetlab and helpful discussions. This project received funding from the FEBS Excellence Award and the European Molecular Biology Laboratory to O.D.

## Author contributions

N.S.Q. and O.D. designed experiments. N.S.Q. cloned, expressed and purified all expressome components, performed all biochemical and single-molecule assays and analyzed all experiments with input from O.D. N.S.Q. and O.D. wrote the manuscript.

