## Supplementary Tables and Figures for "Tracking transcription-translation coupling in real-time"

<u>Content:</u>

Supplementary Tables

Extended Data Figures

Supplementary References

**Supplementary Table 1**: Overview of artificial DNA constructs used in this study

All used DNA sequence have the following common backbone:

RNAP P1 promotor – 5’-immobilization sequence – ribosome binding site – sequence of interest (see below table) – potential binding site for labeled DNA oligo binding to 3’-end of nascent mRNA – T7 terminator sequence

5’-CTGGCAGTTTTAGGCTGATTTGGTTGAATGTTGCGCGGTCAGAAAATTATTTTA AATTTCCTCTTGTCAGGCCGGAATAACTCCCTATAATGCGCCACC – ACTAAAAGA AGAAGAAAGAGAAA – TAGAAGTAATTTTGTTTAAATTTAAGAAGGAGATAT AAA T – SEQUENCE OF INTEREST (flanked by AUG start codon and TAA stop codon) – CCCTATCCCTTATCTTAAC (F1) **OR** AACCACUCCAAUUACAUACACC (F2) – TAATCACACTGGCTCACCTTCGGGTGGGCCTTTCTGCGTTTAT-3’

| DNA construct | Sequence of interest (5’-3’) |
|---|---|
| prNQ215 | ATG TGT GAA AAG AAT GAT TTG GTG GAA GCC CAA AAT AAA TTT GTG AAT ATT CTG TTC GAG ATC CTG GCA CGT TGG AGT TAT GAG TTT CAT CGT CAA TAA |
| prNQ216 | ATG ATT AAA AAT GTG GAA GAT AAT TAT GTT AAA GAT TTG GTG GAA GCC CTG AAT TTT TGT GAG ATT TGG CAT CCG AGC ACC GTG ATG CAG TTT CAA TAA |
| prNQ219 | ATG AAG AAA GAA GTG AAA GTG ATT AAA GAA AAA GTG AAG ATT AAG GAA AAA AAT GTG AAA TTT GAA GTG AAA GAT TTG GTG GAA GCC CAG GTG ACC CAG AAA GAT TTT CTG TGC TTT CAC AGC CGT GTG AAA ATT CTG TAA |
| prNQ291 | ATG TGT GAA AAG AAT AGT GTG AAG GAA GAG GTT AAT GAA ATT AAA ATT ATT AAA AAA ATT ATT GTT ATT ATT GAT AAT TAT GTT AAA AAG AAA GAA GTG AAG GAT GAA GTG TTT AAG AAA GAA GTG AAA GTG ATT AAA GAA AAA GTG AAG ATT AAG GAA AAA AAT GTG AAA GAA GTG AAA GAT TTG GTG GAA GCC CAA AAT AAA TTT GTG AAT ATT CTG TTC GAG ATC CTG GCA CGT TGG AGT TAT GAG TTT CAT CGT CAA TAA |
| prNQ301 | ATG TGT GAA AAG AAT GAA TTA TAT AAT AAG GTT ATT GAA ATG TAT ATG TAT GAT AAA GAA AGT GGT TTA GAA AAA TGG GTG GTT TTG TGG ATT TTA AAA AAG GAA AAG ATG ATT AAA AGT TGT GGT GTT ATG GAA ATT TTA TTA GTG TTA GAT GTA ATT GAA AAT GTA ATT AAT AAA GAG AAG ATG GAG TTA TGG TGT GTT TGG GGG AGT AAT AAA AAG GAT GAA GTG TTT AAG AAA GAA GTG ATG AAA AAT TAT TAT AAA AAG GGA GGA AAA GTA AAT GAA AAA TAT GAT ATT GTG TTA AAT GAA AAA GTT GTA TTA AAA GTG ATT GAG AAA TTG TAT AAG TGG AAA GGT GAA AGT GTA ATT GTG GGG TAT AAA GAT AAG AAA GAA GAA AAT TTG GGT GAA AAG AGT GGA GAT TTA GTT TTA ATG ATT TTG AGT ATT GGT GTG GAT TTG GTG GAA GCC CAA AAT AAA TTT GTG AAT ATT CTG TTC GAG ATC CTG GCA CGT TGG AGT TAT GAG TTT CAT CGT CAA TAA |

**Supplementary Table 2**: Overview of transcribed mRNAs

Below table shows transcribed mRNA sequences. To calculate the intervening mRNA sequence we counted the nucleotides from the ribosome P-site codon till the active site of the RNAP, essentially as defined by Weixlbaumer[1]. Stalling nucleotide is underlined. Note: Zenkin and co-workers counted from A-site codon till the active site (- 3 nt when comparing to us) and Ebright and co-workers counted the codons from A-site till RNA exit channel (- 17 nt when comparing to us).

These are the transcribed RNA sequences:

| DNA construct | Transcribed mRNA sequence (5'-3') | Relevant mRNA lengths |
|---|---|---|
| prNQ215<br>**mRNA-28**<br>(F1) | ACU AAA AGA AGA AGA AAG AGA AAU AGA AGU AAU UUU GUU UAA AUU<br>UAA GAA GGA GAU AUA AAU AUG UGU GAA AAG AAU GAU UUG GUG GAA<br>G**C**C CAA AAU AAA UUU GUG AAU AUU CUG UUC GAG AUC CUG GCA CGU<br>UGG AGU UAU GAG UUU CAU CGU CAA UAA CCC UAU CCC UUA UCU UAA<br>CUA AUC ACA CUG GCU CAC CUU CGG GUG GGC CUU UCU GCG | -91 nt (stalled TEC)<br>-219 nt (Full-length)<br>-**28** nt between P-site and active site at start<br>-156 nt between P-site and active site at TC end |
| prNQ216<br>**mRNA-46**<br>(F1) | ACU AAA AGA AGA AGA AAG AGA AAU AGA AGU AAU UUU GUU UAA AUU<br>UAA GAA GGA GAU AUA AAU AUG AUU AAA AAU GUG GAA GAU AAU UAU<br>GUU AAA GAU UUG GUG GAA G**C**C CUG AAU UUU UGU GAG AUU UGG CAU<br>CCG AGC ACC GUG AUG CAG UUU CAA UAA CCC UAU CCC UUA UCU UAA<br>CUA AUC ACA CUG GCU CAC CUU CGG GUG GGC CUU UCU GCG | -109 nt (stalled TEC)<br>-219 nt (Full-length)<br>-**46** nt between P-site and active site at start<br>-71 nt intervening to P1<br>-106 nt intervening to P2<br>-156 nt between P-site and active site at TC end |
| prNQ219<br>**mRNA-85**<br>(F1) | ACU AAA AGA AGA AGA AAG AGA AAU AGA AGU AAU UUU GUU UAA<br>AUU UAA GAA GGA GAU AUA AAU AUG AAG AAA GAA GUG AAA GUG AUU<br>AAA GAA AAA GUG AAG AUU AAG GAA AAA AAU GUG AAA UUU GAA GUG<br>AAA GAU UUG GUG GAA G**C**C CAG GUG ACC CAG AAA GAU UUU CUG UGC<br>UUU CAC AGC CGU GUG AAA AUU CUG UAA CCC UAU CCC UUA UCU UAA<br>CUA AUC ACA CUG GCU CAC CUU CGG GUG GGC CUU UCU GCG | -148 nt (stalled TEC)<br>-261 nt (Full-length)<br>-**85** nt between P-site and active site at TC end<br>-198 nt between P-site and active site at TC end |
| prNQ291<br>**mRNA-193**<br>(F2) | ACU AAA AGA AGA AGA AAG AGA AAU AGA AGU AAU UUU GUU UAA AUU<br>UAA GAA GGA GAU AUA AAU AUG UGU GAA AAG AAU AGU GUG AAG GAA<br>GAG GUU AAU GAA AUU AAA AUU AUU AAA AAA AUU AUU GUU AUU AUU<br>GAU AAU UAU GUU AAA AAG AAA GAA GUG AAG GAU GAA GUG UUU AAG<br>AAA GAA GUG AAA GUG AUU AAA GAA AAA GUG AAG AUU AAG GAA AAA<br>AAU GUG AAA GAA GUG AAA GAU UUG GUG GAA G**C**C CAA AAU AAA UUU<br>GUG AAU AUU CUG UUC GAG AUC CUG GCA CGU UGG AGU UAU GAG UUU<br>CAU CGU CAA UAA AAC CAC UCC AAU UAC AUA CAC CUA AUC ACA CUG<br>GCU CAC CUU CGG GUG GGC CUU UCU GCG | -256 nt (stalled TEC)<br>-387 nt (Full-length)<br>-**193** nt between P-site and active site at start<br>-324 nt between P-site and active site at start |
| prNQ301<br>**mRNA-457**<br>(F2) | ACU AAA AGA AGA AGA AAG AGA AAU AGA AGU AAU UUU GUU UAA AUU<br>UAA GAA GGA GAU AUA AAU AUG UGU GAA AAG AAU GAA UUA UAU AAU<br>AAG GUU AUU GAA AUG UAU AUG UAU GAU AAA GAA AGU GGU UUA GAA<br>AAA UGG GUG GUU UUG UGG AUU UUA AAA AAG GAA AAG AUG AUU AAA<br>AGU UGU GGU GUU AUG GAA AUU UUA UUA GUG UUA GAU GUA AUU GAA<br>AAU GUA AUU AAU AAA GAG AAG AUG GAG UUA UGG UGU GUU UGG GGG<br>AGU AAU AAA AAG GAU GAA GUG UUU AAG AAA GAA GUG AUG AAA AAU<br>UAU UAU AAA AAG GGA GGA AAA GUA AAU GAA AAA UAU GAU AUU GUG<br>UUA AAU GAA AAA GUU GUA UUA AAA GUG AUU GAG AAA UUG UAU AAG<br>UGG AAA GGU GAA AGU GUA AUU GUG GGG UAU AAA GAU AAG AAA GAA<br>GAA AAU UUG GGU GAA AAG AGU GGA GAU UUA GUU UUA AUG AUU UUG<br>AGU AUU GGU GUG GAU UUG GUG GAA G**C**C CAA AAU AAA UUU GUG AAU<br>AUU CUG UUC GAG AUC CUG GCA CGU UGG AGU UAU GAG UUU CAU CGU<br>CAA UAA AAC CAC UCC AAU UAC AUA CAC CUA AUC ACA CUG GCU CAC<br>CUU CGG GUG GGC CUU UCU GCG | -520 nt (stall)<br>-651 nt (Full-length)<br>-**457** nt between P-site and active site at start<br>-588 nt between P-site and active site at start |

**Supplementary Table 3**: Overview of DNA oligonucleotides used for biochemical and/or single-molecule assays.

All DNA oligonucleotides were purchased from IDT or biomers.

| Oligonucleotide ID | DNA sequence (5'-3') |
|---|---|
| p0030_ab_fw | TCACGAAAGCTGAGTAGTCACGAGTCTTCT/idSp/CTGGCAGTTTTAGGCTGATTTGG |
| p0075_ab_bw | CCTTAATCATACTCACCAAATTACCATCCC/idSp/ATAAACGCAGAAAGGCCCAC |
| p0088_2xCy3.5 | Cy3p5-GGGATGGTAATTTGG[dT-Cy3p5]GAGTATGATTAAGG |
| p0109-biotin | 5BiotinTEG/TTATCCGCTCACAATTCCACA |
| prNQ087-Cy3 | GGGAGATCAGGATA/3Cy3Sp/ |
| prNQ088-Cy5 | GAGGCCGAGAAGTG/3Cy5Sp/ |
| prNQ127 | TTTCTCTTTCTTCTTCTTTTAGTTGTGGAATTGTGAGCGGATAA |
| prNQ302 | GAGGCCGAGAAGTGAAAAACCACTAGTCCACCGCAGCCC |
| prNQ303 | TACCTATGGATCCATATCTGGGGCTGCGGTGGACTAGTGG |
| prNQ304 | CAGATATGGATCCATAGGTATGTGGAATTGTGAGCGGATAA |

/idSp/ denotes abasic site.
[dT-Cy3p5] is an internal modification labeled on the nucleobase of deoxythymidine.
5BiotinTEG denotes Biotin-TEG attached to 5'-end
/3Cy3sp/ denotes Cy3 label attached to 3'-end
/3Cy5sp/ denotes Cy5 label attached to 3'-end

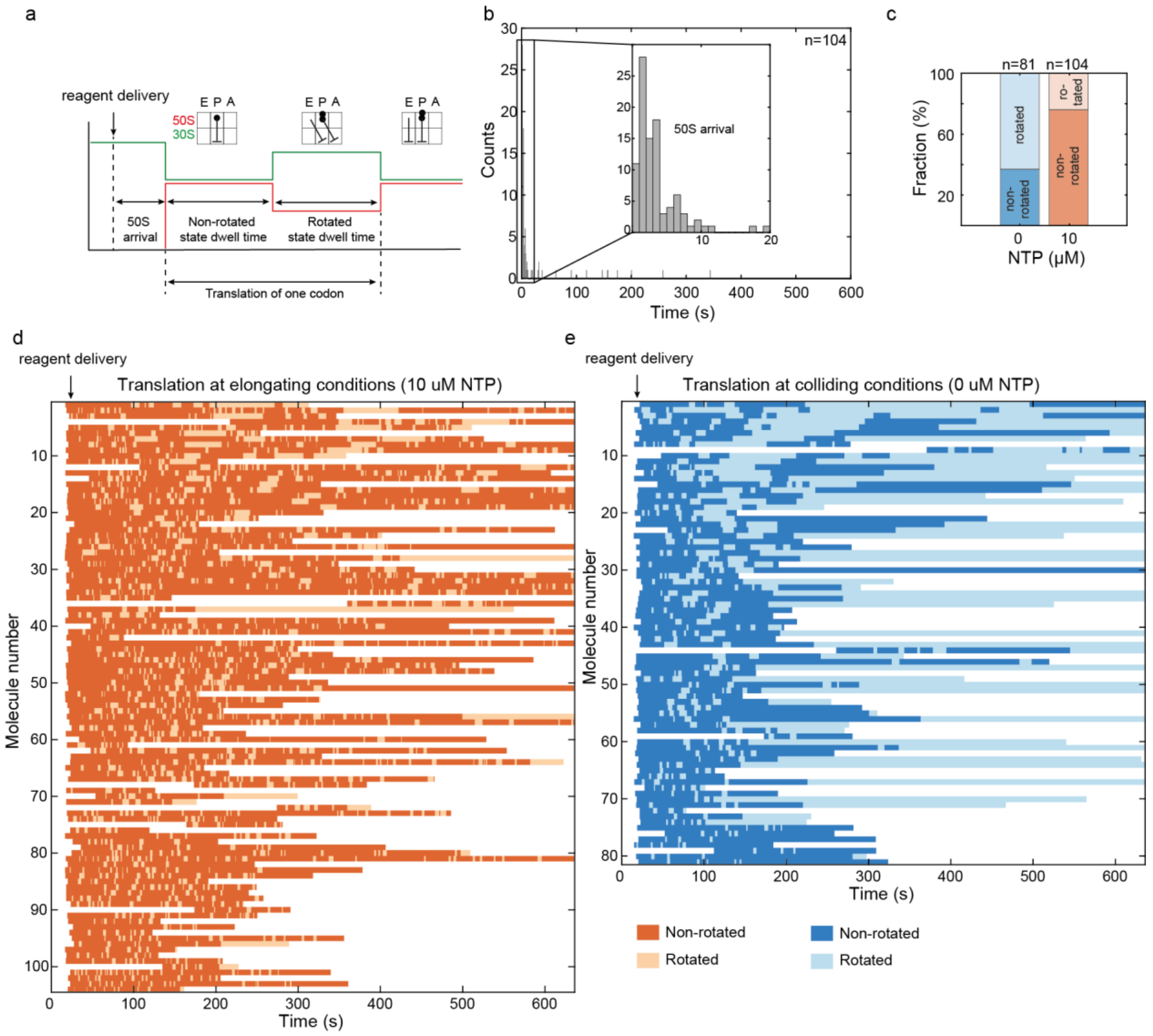


**Extended Data Figure 1**. **a,** Scheme for data evaluation. **b,** 50S arrival time to the mRNA-46 stalled TEC loaded with 30S PIC. Data was acquired with 50 nM 50S-Cy5, 2 µM IF2a, 150 nM aa-tRNA, 450 nM EF-Tu, 300 nM EF-Ts, 50 nM EF-G at 21°C. The time of 50S arrival to the expressome is plotted as histogram. Inset shows zoom of the first 20 s after reagent delivery. Reagent delivery time is at t=0 s. **c,** Fraction of molecules ending in rotated state or non-rotated state for colliding (0 µM NTP) or elongating (10 µM NTP, each) conditions. Last state before photobleaching was evaluated. A total of 63% of traces in colliding conditions stall in rotated state. A total of 24% of traces in elongating conditions stall in rotated state. **d, e,** Stack of single-molecule traces for elongating (left) and colliding (right) conditions. Each row represents a single transcription-translation complex. Non-rotated states are displayed in dark orange (elongating) or dark blue (colliding). The respective rotated states are shown in light orange or light blue.

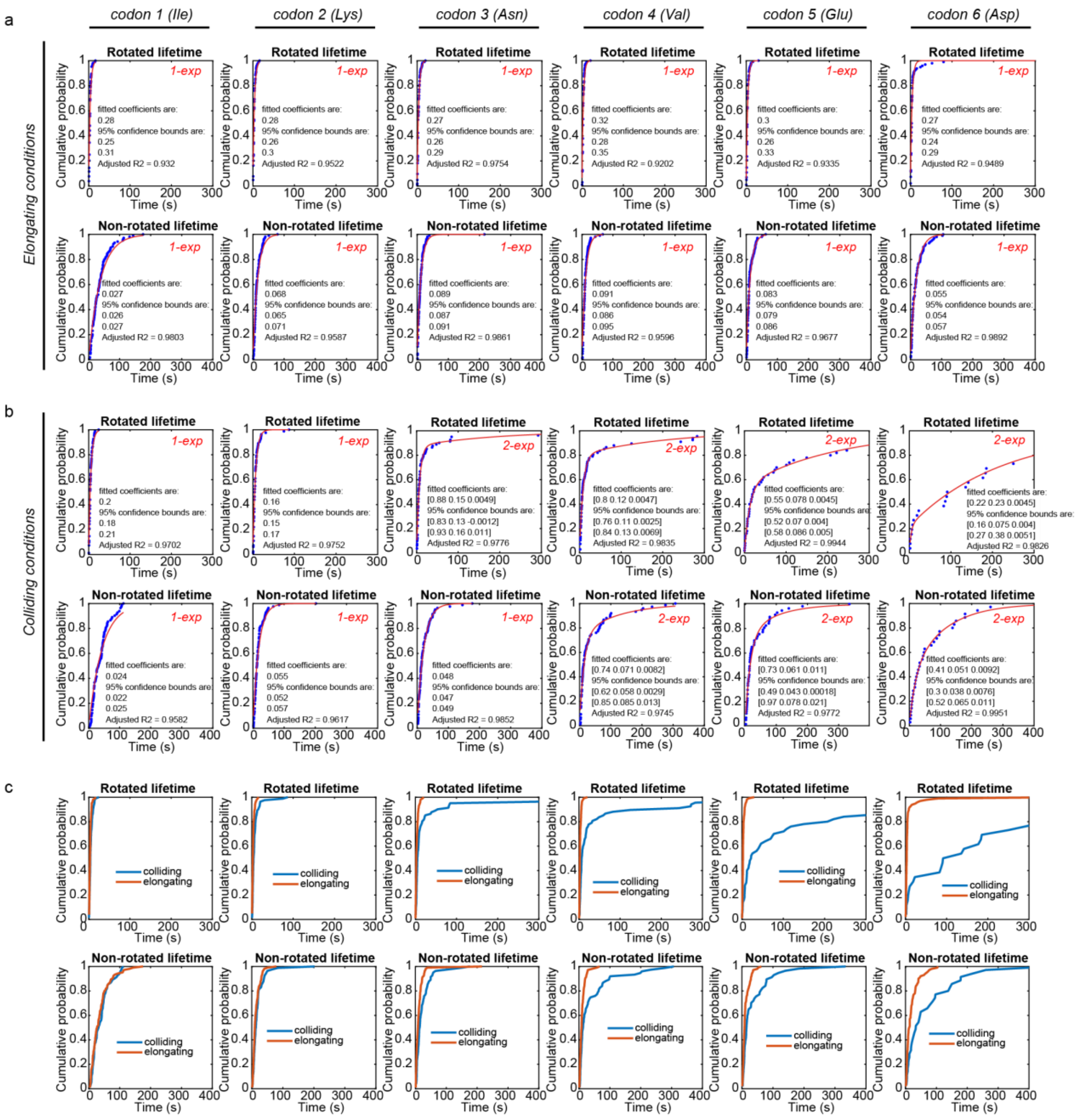


**Extended Data Figure 2**. Single and double exponential fitting of non-rotated and rotated state dwell-times for the first 6 amino acids during elongating and colliding conditions. **a,** Elongating conditions can be fitted with a single exponential function y=1-exp(-b*t), with given coefficient [b] representing the mean transition rate for the respective state. **b,** In case of colliding conditions, only the first two codon translations can be fitted with a single exponential function. Amino acids 3-6 (rotated state) and amino acids 4-6 (non-rotated state) are fitted with a double exponential function y=1-a1*exp(-b1*t)-(1-a1)*exp(-b2*t). Fitted coefficients are given as [a1, b1, b2]. **c**, Cumulative probability of colliding (blue) and elongating (orange) conditions are overlayed.

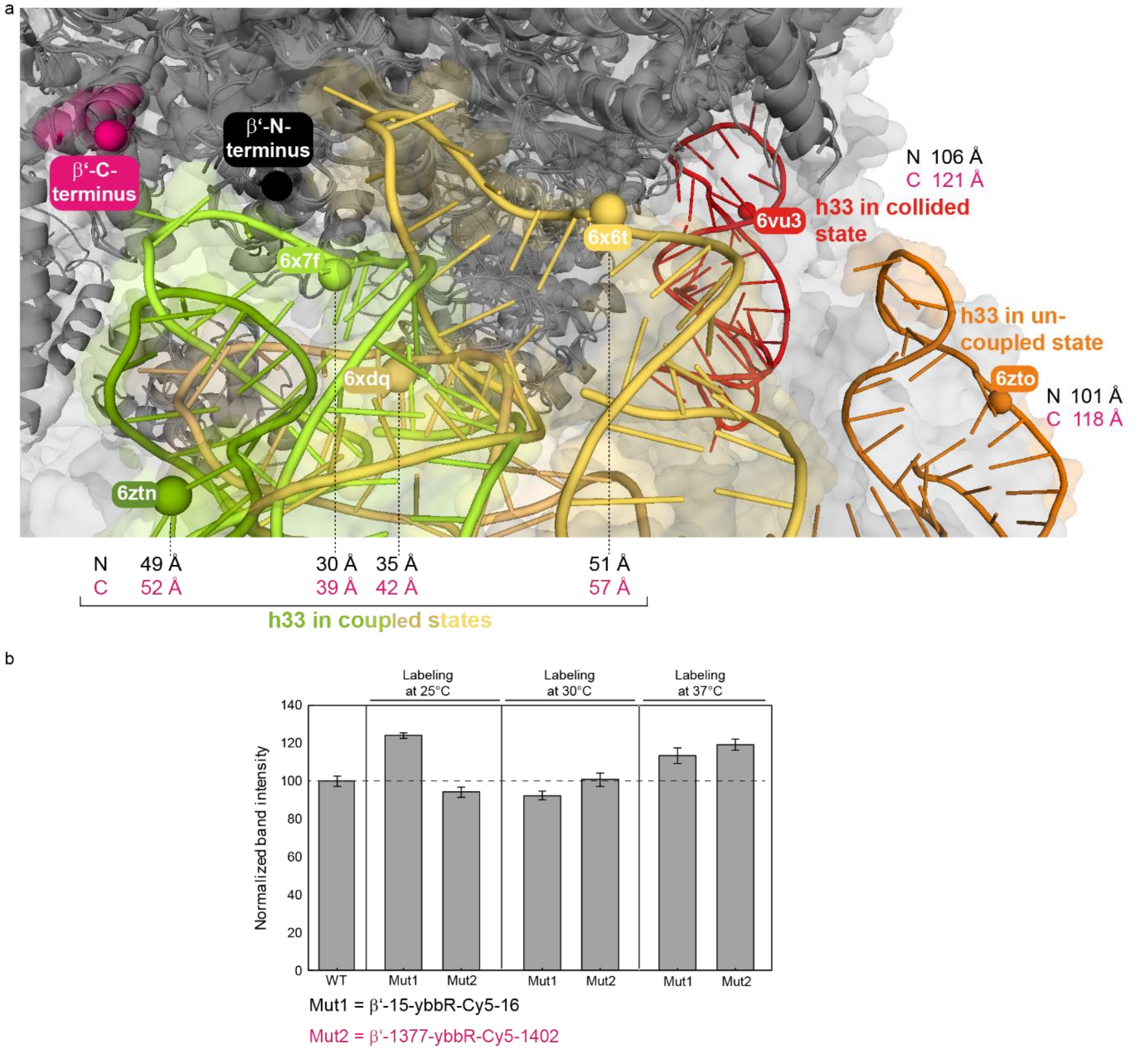


**Extended Data Figure 3**. **a**, Overlay of expressome structures (6ztn, 6x7f, 6xdq, 6x6t, 6vu3, 6zto)[1,2] showing the variation of FRET distances between the ribosomal h33a (16S rRNA) and RNAP β' (Nter=black and Cter=pink) labeling sites. Structures were aligned on the RNAP. Helix 33 is color coded according to PDB ID and displayed as cartoon representation. Labeling sites on RNAP and 16S rRNA are indicated as spheres. In all displayed coupled states, the labeling sites are in FRET distance, whereas in the collided state and uncoupled state, they are too far to be detected by FRET (>100 Å). **b,** RNAP-Cy5 activity test using single-round transcription assays. Area of total RNA was integrated and normalized to WT RNAP. Introduction of the ybbR-peptide tag as well as the Cy5 label do not significantly affect RNAP activity.

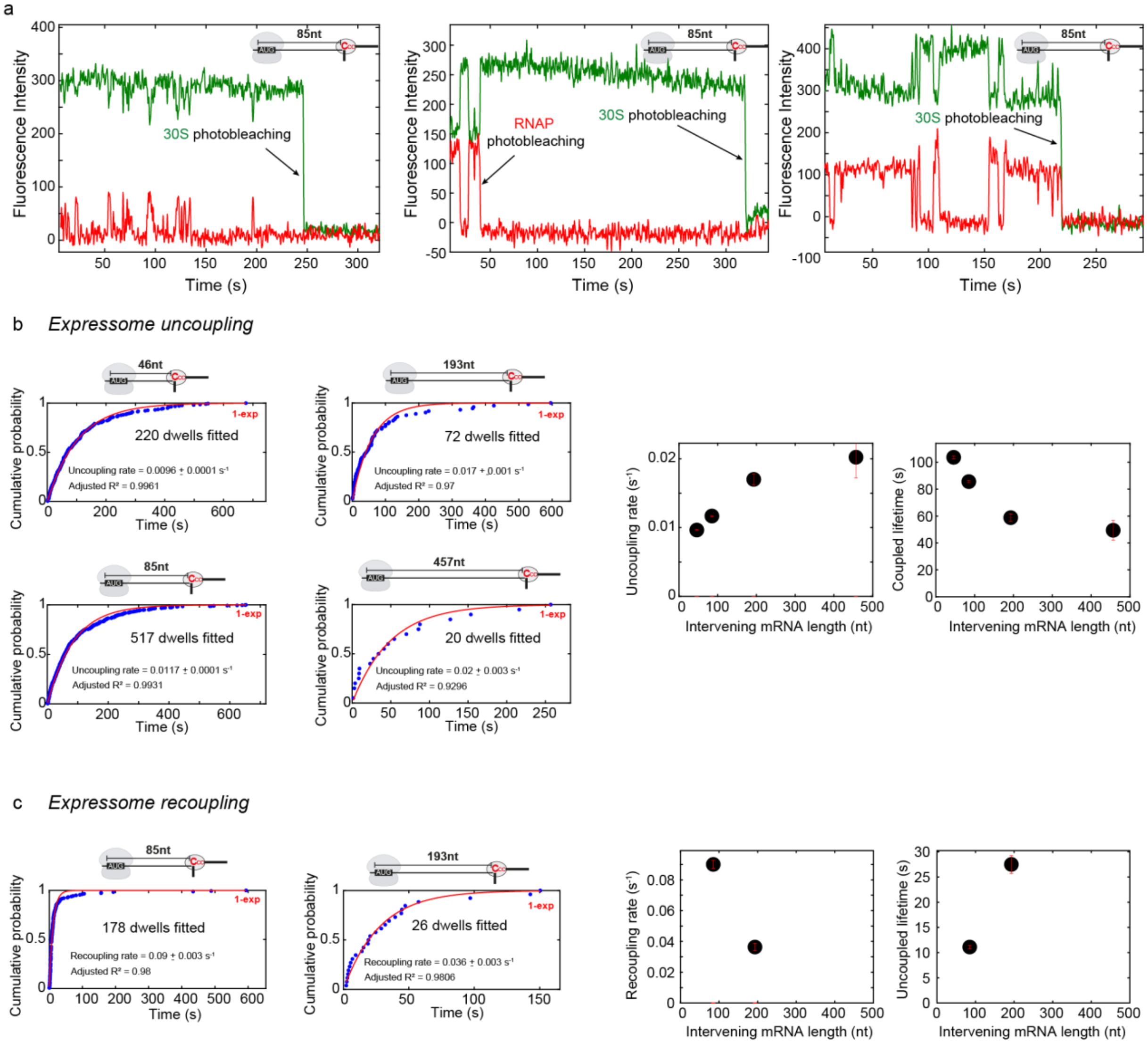


**Extended Data Figure 4**. **a,** Representative single-molecule traces (mRNA-85) displaying dynamically coupled expressome molecules acquired as equilibrium experiments (no reagent delivery). **b, c,** Expressome uncoupling (**b**) and recoupling (**c**) dynamics in dependence of mRNA length. The coupling (periods of 30S-Cy3/RNAP-Cy5 FRET) or uncoupling (periods of no Cy3-Cy5 FRET) signals were evaluated and the dwells were fitted with a single (y=1-exp(-b*t)) or double (y=1-a1*exp(-b1*t)-(1-a1)*exp(-b2*t)) exponential equation. All dwells, included photobleaching-limited ones, were included in the fitting.

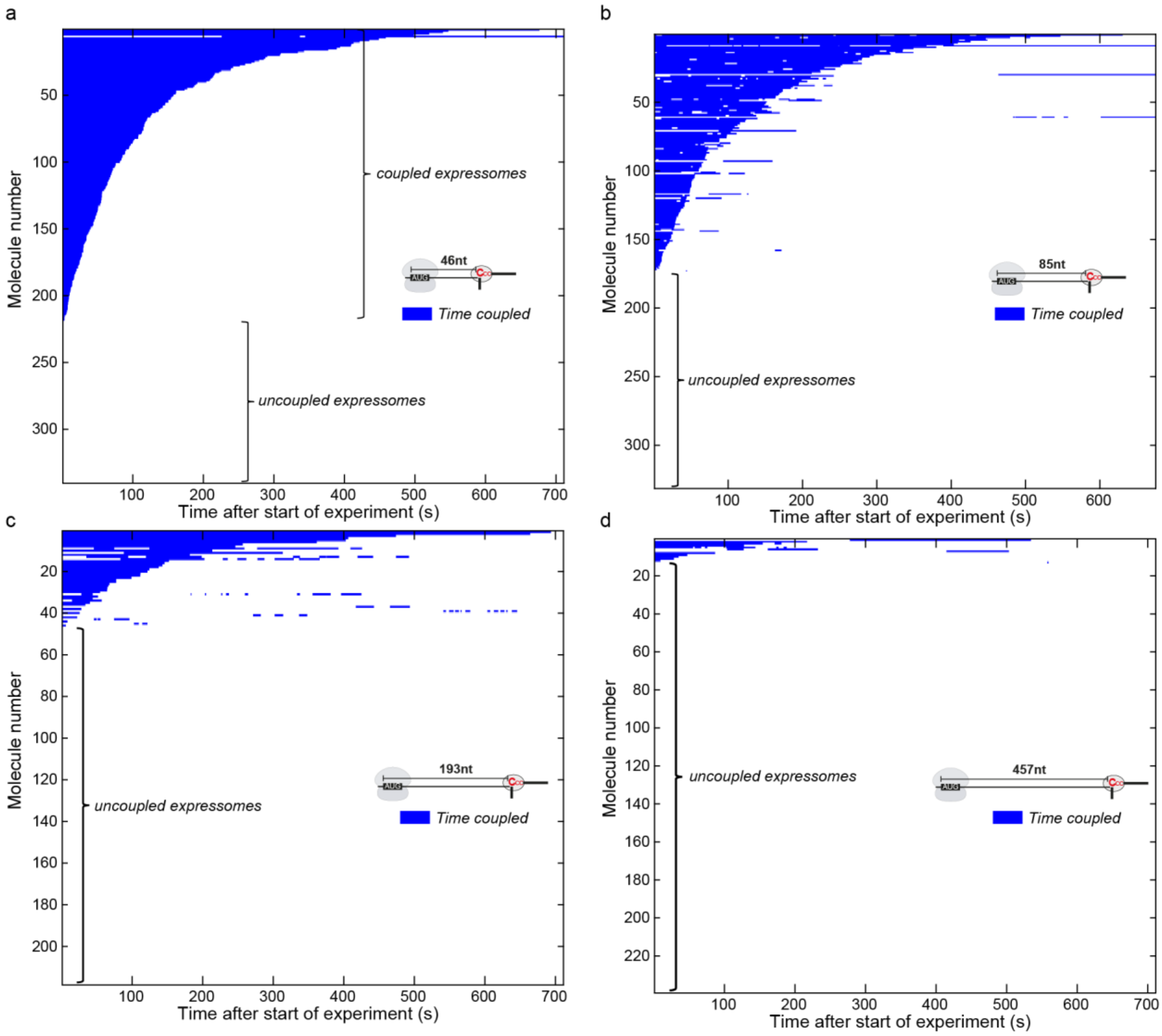


**Extended Data Figure 5**. Stack of raw single-molecule traces for **a**, mRNA-46, **b,** mRNA-85, **c,** mRNA-193, **d,** mRNA-457 in absence of transcription and translation elongation. Each row represents a single transcription-translation complex. Coupled states (characterized by 30S-Cy3/RNAP-Cy5 FRET signal) are shown in blue. Traces are sorted by the total time for which coupling can be detected. White spaces in between coupling events represent uncoupled expressomes. The fraction of uncoupled expressomes increases with mRNA length. In case of the mRNA-46 expressome, we do not detect any uncoupling events and therefore, the apparent coupled state lifetime is limited by photobleaching. Traces without any coupled states during the entire experimental time are represented as empty traces (white) at the bottom of each plot.

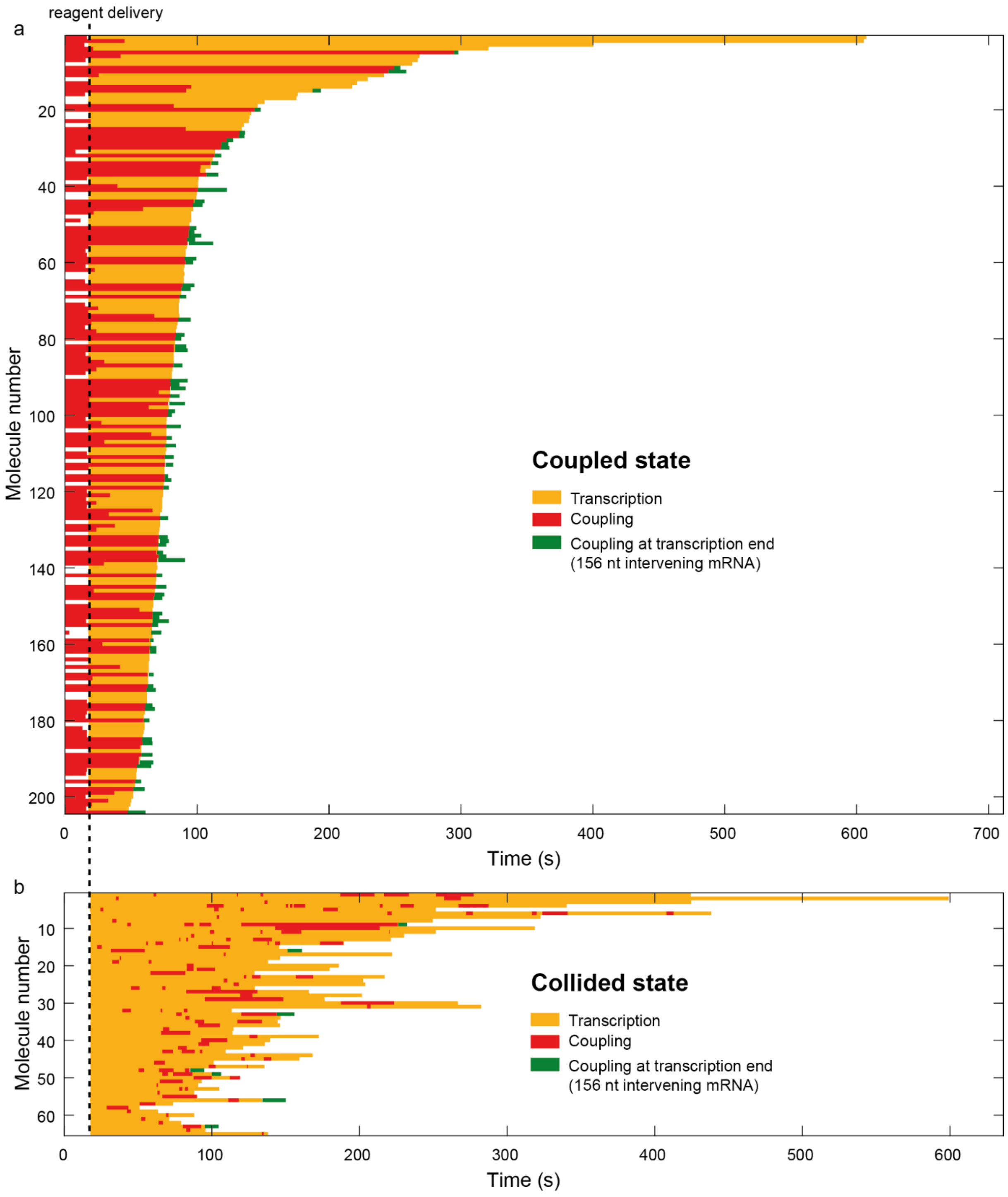


**Extended Data Figure 6**. Stack of single-molecule traces for transcriptions out of coupled (**a**) or collided (**b**) state. Each row represents a single transcription-translation complex. Experiment start was triggered by delivering 50 µM NTPs to the immobilized and stalled expressome molecules in absence of Nus factors. Transcriptions are depicted in yellow-orange, coupling events are shown in red and traces with coupling at transcription end are marked in green. Note: For the coupled state construct, all single-expressome molecules are shown. For the collided state construct, only single-expressome molecules are shown that showed coupling at least once during subsequent transcription elongation. A large portion of collided expressomes failed to establish coupling completely during subsequent transcription elongation. Therefore, the difference in coupling efficiency following transcription out of a coupled versus collided expressome is even larger as it appears from comparing above clusters.

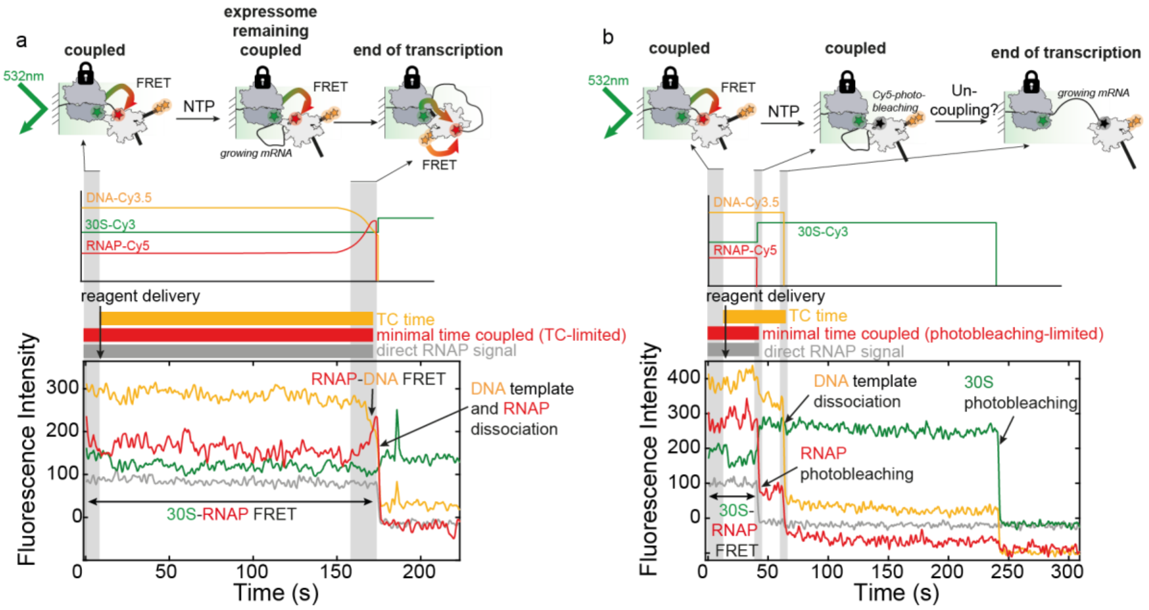


**Extended Data Figure 7**. Representative traces for transcription out of coupled state. Data was acquired with alternative laser excitation at wavelengths of 532nm and 638nm. The reaction was started with delivery of 50 µM NTP (each). The 70S ribosome was kept stalled on the RBS. **a**, Illustration of single-molecule trace, where both machines remain coupled throughout the complete transcription reaction. The steady increase in Cy3.5-Cy5 FRET efficiency towards the end of transcription (at ~160-170 s) directly shows active transcription elongation while both machines are coupled containing an intervening mRNA length of 156 nt. **b**, Single-molecule trace with photobleaching of the RNAP before transcription is completed. Time-evolution of coupling cannot be tracked by 30S-Cy3 and RNAP-Cy5 FRET anymore. Moreover, the expressome uncouples after RNAP-photobleaching and before transcription end, as also no 30S-Cy3 to DNA-Cy3.5 FRET is detected (see **Fig. 4a**).

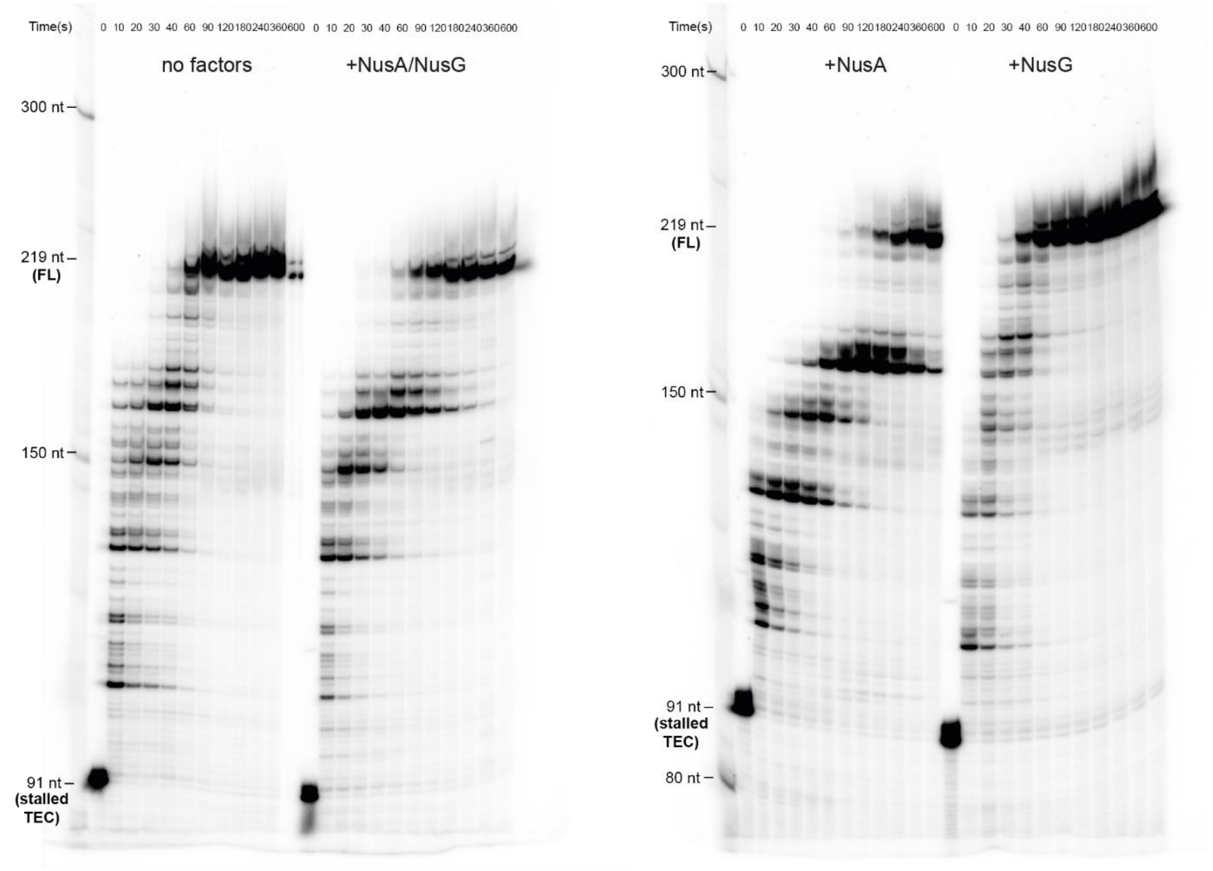


**Extended Data Figure 8**. Single-round transcription assays with assembled 70S collided expressome using prNQ215 DNA template. Stalled transcription elongation complex (stalled TEC) was formed with 50 nM DNA template, 200 nM *E. Coli* RNAP, 100 µM ACU trinucleotide, 5 µM GTP and 5 µM ATP (+ 150 nM $^{32}$P α-ATP) halting RNAP initially at U24 to prevent loading of multiple RNAPs. Then RNAP was walked to desired stalling site by addition of 10 µM UTP and simultaneous addition of 10µg/mL rifampicin (to prevent transcription re-initiation). 70S PIC was formed on the stalled TEC in presence of 2 µM IF2a, 1 µM fmet-tRNA$^{fmet}$ and 4 mM GTP. This stalled expressome was chased in presence or absence of NusA and/or NusG with 50 µM NTPs (each) at room temperature and per condition time points were taken at 0, 10, 20, 30, 40, 60, 90, 120, 180, 240, 360 and 600 seconds. Stalled expressome band (91 nt) was immediately chased after NTP addition (<10 s).

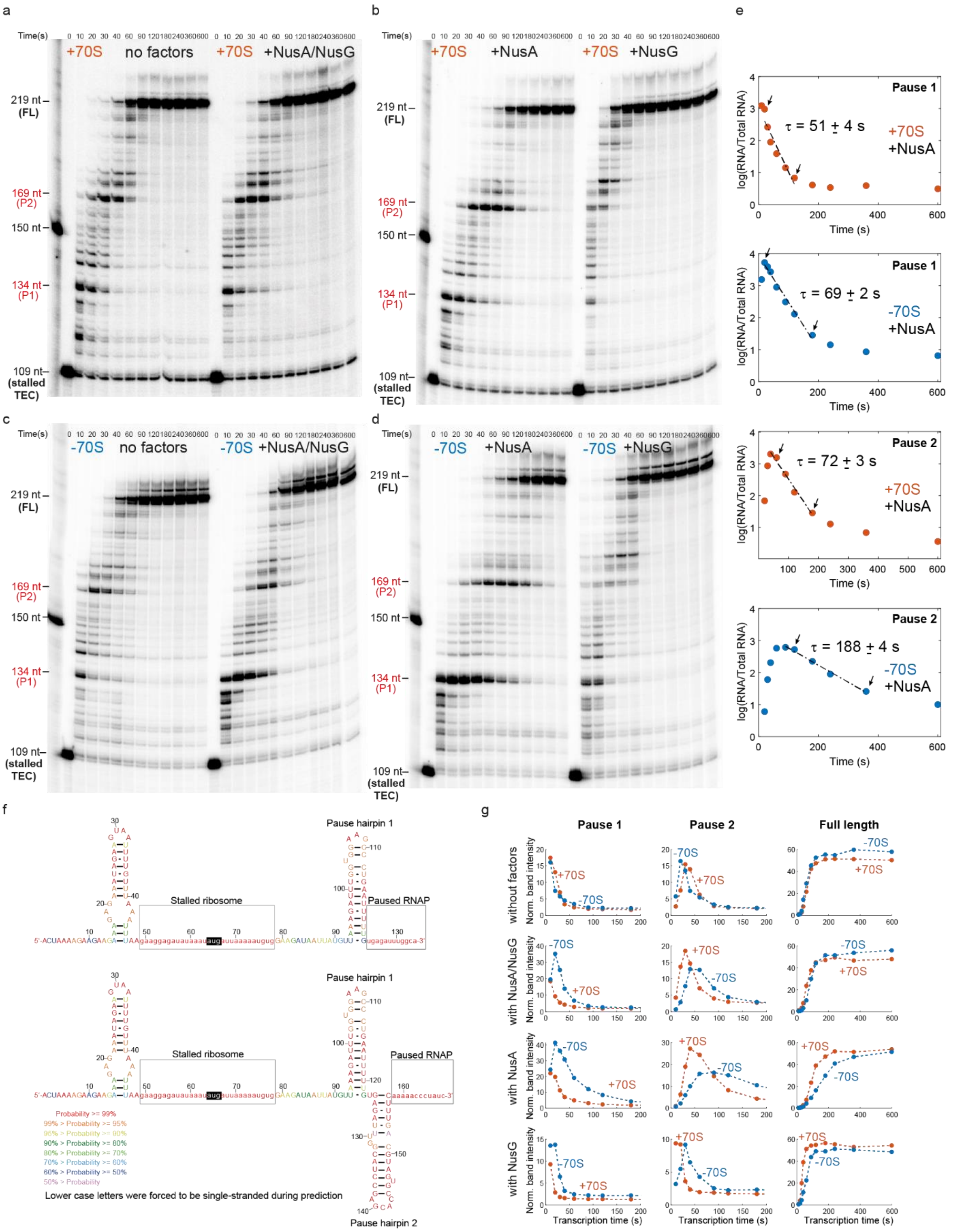


**Extended Data Figure 9**. **a, b, c, d,** Single-round transcription assays with coupled expressome using the prNQ216 DNA template. Stalled transcription elongation complex (stalled TEC) was formed with 50 nM DNA template, 200 nM *E. Coli* RNAP, 100 µM ACU trinucleotide, 5 µM GTP and 5 µM ATP (+ 300 nM $^{32}$P α-ATP) halting RNAP initially at U24 to prevent loading of multiple RNAPs. Then RNAP was walked to desired stalling site by addition of 10 µM UTP and simultaneous addition of 10 µg/mL rifampicin (to prevent re-initiation). The 70S PIC was formed on the stalled TEC in presence of 2 µM IF2a, 1 µM fmet-tRNA$^{fmet}$ and 4 mM GTP using 250 nM C68-30S and 500 nM ZS22-50S mutants[3,4]. h101 of the 50S mutant was pre-annealed with a biotin-oligonucleotide and the formed expressomes were purified using streptavidin beads to enrich for fully assembled transcription-translation complexes. The purified stalled expressome (+70S) or the stalled transcription complex alone (-70S) were chased in presence or absence of Nus factors (w/o factors, w/ NusA/NusG, w/ NusA only, w/ NusG only; 1 µM final concentration for each Nus factor) with 50 µM NTP (each) at room temperature. For each condition, time points were taken at 0, 10, 20, 30, 40, 60, 90, 120, 180, 240, 360 and 600 seconds. **e**, Pause escape lifetimes for pause 1 and pause 2 in presence of NusA and in presence (orange) or absence (blue) of ribosome. Natural logarithm of normalized band intensities (P/T) was plotted as function of time and pause escape lifetimes were fitted with a linear fit function (y=m*x+b, with m being the rate constant)[5]. Data range that was used for fitting is indicated with arrows. **f,** Secondary structure prediction of the nascent mRNA (mRNA-46) using the RNA structure web server (https://rna.urmc.rochester.edu/RNAstructureWeb/). Top prediction shows secondary structure at pause site 1 (134 nt) and bottom prediction shows secondary structure at pause site 2 (169 nt). Secondary structure was predicted using default settings on the website for 21 °C, forcing the ribosome binding site (as it is masked by the ribosome) and 12 nt (reference [6]) upstream from 3'-end of the nascent mRNA (as they are masked by the paused RNAP) to be single-stranded. Color code corresponds to probability of base-pair formation. Position of the ribosome and the RNAP are indicated. Start codon (AUG) is highlighted with a black box. **g**, Normalized band intensities (P/T) are displayed as a function of time. Bands for pause 1, pause 2 and full-length (FL) RNA were integrated and divided by the total RNA per lane[5].

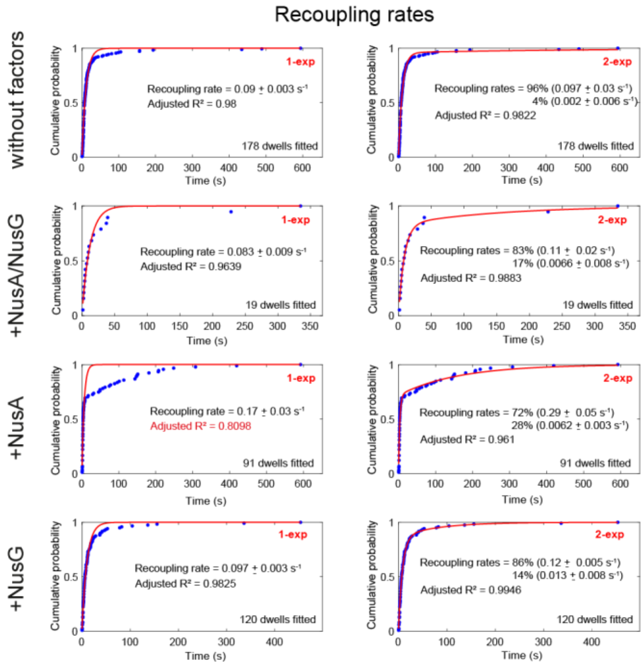


**Extended Data Fig. 10.** Recoupling dynamics for mRNA-85 in absence of transcription and translation elongation and in presence and absence of Nus factors at 1 µM (each). The data was acquired as equilibrium experiments (no reagent delivery) monitoring the coupling dynamics of transcription-translation complexes with alternative laser excitation over time. The recoupling dwell times were fitted with single (y=1-exp(-b*t)) or double (y=1-a1*exp(-b1*t)-(1-a1)*exp(-b2*t)) exponential equations.
